# Toward a genetic signature of resistance to activity-based anorexia in striatal projecting cortical neurons

**DOI:** 10.1101/2025.03.09.642272

**Authors:** K Huang, MA Magateshvaren Saras, K Conn, E Greaves, F Reed, S Tyagi, H Munguba, CJ Foldi

## Abstract

**Objective:** Converging evidence from neuroimaging studies and genome-wide association study (GWAS) suggests the involvement of prefrontal cortex (PFC) and striatum dysfunction in the pathophysiology of anorexia nervosa (AN). However, identifying the causal role of circuit-specific genes in the development of AN-like phenotype remains challenging and requires the combination of novel molecular tools and preclinical models.

**Methods:** We used the activity-based anorexia (ABA) rat model in combination with a novel viral-based translating ribosome affinity purification (TRAP) technique to identify transcriptional differences within a specific neural pathway that we have previously demonstrated to mediate pathological weight loss in ABA rats (i.e. medial prefrontal cortex neurons that project to the nucleus accumbens shell). We compared actively transcribed genes in rats susceptible to weight loss to the subpopulation of rats resistant to weight loss under the same experimental conditions.

**Results:** We reveal 1424 differentially expressed genes between Susceptible and Resistant rats, highlighting important transcriptional changes associated with ABA within this pathway. The changes observed were independent of current calorie deficit and associated with metabolic, mitochondrial and neural functions. Further, we show that genes upregulated in Resistant rats were involved in mitochondrial function, while downregulated genes were associated with cytoskeletal, postsynaptic and axonal functions, supporting the hypothesis that hyperexcitability of cortico-striatal circuit function is a critical mediator of pathological weight loss in ABA.

**Discussion:** These findings represent an essential first step in understanding how circuit-specific gene expression patterns may contribute to susceptibility to ABA and provide potential molecular targets for manipulation in this animal model of AN.

**Public Significance:** This study identifies specific brain gene activity patterns that may explain why some individuals are more vulnerable to extreme weight loss, as seen in anorexia nervosa. Using an advanced molecular technique in a well-established animal model, key differences in a neural pathway linked to cognitive control were observed. These findings pave the way for more targeted treatments that could prevent or reverse this dangerous condition.

**Summary:**

- Transcriptional differences within medial prefrontal cortex to nucleus accumbens shell (mPFC–AcbSh) neurons distinguish rats highly susceptible to activity-based anorexia (ABA) from those resistant to pathological weight loss.
- Resistant rats showed upregulation of mitochondrial function genes, while Susceptible rats had upregulation of genes related to synaptic structure and signalling, implicating excitability of this circuit in driving maladaptive weight loss behaviour.
- Transcriptomic changes align with human genome-wide association study (GWAS) findings and support links between anorexia nervosa and both metabolic and psychiatric comorbidities.
- The study highlights potential molecular targets for future gene manipulation and therapeutic intervention. It also provides a foundation for creating refined genetic animal models that integrate multiple AN-associated variants to better reflect the polygenic nature of the disorder.

## Introduction

Anorexia nervosa (AN) is a complex and serious illness characterised by a low body-mass index, behavioural disturbances such as inflexible thinking and compulsive behaviours, and higher mortality rates than most psychiatric disorders (Chesney, Goodwin, & Fazel, 2014). Available treatments have low rates of clinical success, in part, because renourishment protocols are not efficacious in addressing the deleterious drive to exacerbate dangerously low body weights (Watson & Bulik, 2013). Converging evidence from human neuroimaging studies and preclinical models suggest the involvement of altered neural circuit function, particularly aberrant activity within the prefrontal cortex (PFC) and striatum regions of the brain (Cha et al., 2016; Ehrlich et al., 2015; C. J. Foldi, Milton, & Oldfield, 2017; Milton et al., 2021). Interestingly, the largest genome-wide association study (GWAS) conducted to date (including ∼17K cases and 55K controls) (Watson et al., 2019) suggests that genes associated with AN are enriched in similar brain regions, namely PFC and striatum (Song, Wang, Yu, & Lin, 2021), activity in which is associated with food reward valuation and cognitive flexibility (Cox & Witten, 2019; Joshi, Schott, la Fleur, & Barrot, 2022). Although GWAS findings are informative, identifying strong hypotheses about their connections to specific genes, or how these specific genes are causal to changes in brain function or behaviour is not straightforward. Using a sophisticated complement of analytical tools including expression quantitative trait loci (eQTL) and chromosome conformation capture (Hi-C) interactions, 58 specific genes were identified to be significantly dysregulated in AN cases, with the clearest evidence for a role in the aetiology of AN for *Cadm1*, *Mgmt*, *Focp1* and *Ptbp2* (Watson et al., 2019), genes that are involved in cellular processes like receptor binding, DNA repair and cell-type specific gene transcription during development. However, it is important to recognise that by definition these findings are associations and may reflect variants and genes with no direct biological relevance to the causes of AN. Moreover, understanding the genetic predisposition of any neuropsychiatric disease is insufficient for informing novel prevention or treatment strategies if it does not pinpoint the brain regions or circuits where those genes have the greatest impact. Insight into the causal role of circuit-specific genes in the development and/or maintenance of AN- like phenotypes is now possible with novel molecular tools in combination with preclinical models.

Activity-based anorexia (ABA) is a biobehavioural rodent model that recapitulates many of the behavioural phenotypes of AN in humans, including voluntary food restriction and paradoxical hyperactivity in states of negative energy balance that leads to rapid reductions in body weight. Accordingly, the ABA model has been used for decades to gain mechanistic insight into the biological bases for AN, including predisposing factors (Barbarich-Marsteller et al., 2013; Carrera, Gutiérrez, & Boakes, 2006; Hancock & Grant, 2009; Milton, Patton, O’Keeffe, Oldfield, & Foldi, 2022). The observation that when exposed to the same environmental conditions of ABA, both wild-type adolescent rats (Milton, Oldfield, & Foldi, 2018) and mice (Beeler & Burghardt, 2021) consistently split into ‘Susceptible’ and ‘Resistant’ phenotypes makes this a particularly powerful model for understanding what differentiates individuals that go on to develop pathological weight loss and compulsive exercise from those that do not. Therefore, the ABA model could provide a valuable tool to dissect the causal involvement of genes associated with AN, but thus far its use has not shed sufficient light on the genetic determinants of AN (Foldi, 2024). We have previously shown that suppressing activity within a cortico-striatal neural circuit that extends between the medial prefrontal cortex (mPFC) and nucleus accumbens shell (AcbSh) both prevents weight loss in ABA and improves cognitive flexibility in female rats (Milton et al., 2021). This circuit is important in the context of AN-like behaviours because of its role in the suppression of learned stimulus-response behaviour (i.e. behaviour that is influenced by the consequences of actions) by regulating sensitivity to punishment (Piantadosi, Yeates, & Floresco, 2020). Moreover, dysfunction in this circuit is reflected in the analogous human brain regions (PFC, ventral striatum) in individuals with AN, which may increase throughout the course of illness (Fladung, Schulze, Schöll, Bauer, & Grön, 2013) and persist after weight recovery (Ehrlich et al., 2015). A caveat in interpreting our circuit-level findings arises from a subsequent study in which we examined flexible learning and susceptibility to ABA in the same (rat) subjects. Here, we showed that in fact *better* performance on the cognitive task predicted weight loss in ABA (Huang et al., 2023) suggesting a more nuanced role for cortico-striatal circuitry and reinforcement learning in the development of ABA and possibly the prodromal symptoms of AN in humans.

In order to interpret how disrupted neurochemistry and/or neural circuit function gives rise to the specific pathology of AN, the molecular phenotype that determines the functional output of the neural circuits involved needs to be identified. Traditional assessment of the genetic contributions to AN using targeted genetic knockout/down in animal models have yielded insufficient evidence to date for a molecular signature of AN (Scharner & Stengel, 2021), perhaps because of compensatory mechanisms that come into play when a gene construct is manipulated within an entire organism or brain region. It is also important in the context of a condition like AN, for which very little is known about the underlying genetic drivers, to perform an unbiased screen of gene expression within specific circuits of ABA rats, in order to reveal novel mechanisms. With this in mind, we utilised translating ribosome affinity purification (TRAP) technology, which allows the synthesis of molecular and neuroanatomical information by immunoprecipitation of translating mRNAs from any population of neurons that express enhanced green fluorescent protein (eGFP). While this technique has been traditionally used to identify cell type-specific genetic profiles (Heiman, Kulicke, Fenster, Greengard, & Heintz, 2014) more recently it has been used effectively to differentiate genetic signatures of addiction-like behaviour to non-addicted controls (Kawa, Hashimoto, Beutler, Guizzetti, & Wolf, 2025). Thus, in order identify differentially regulated genes *within a specific neural pathway* that modulates weight loss in ABA rats, we employed a dual viral approach, in which coincident injection of a retrogradely transporting Cre-expressing virus and a Cre-dependent TRAP construct (eGFP-tagged ribosome protein L10a; EGFPL10a) were injected into the AcbSh and mPFC, respectively. This allows eGFP-tagging of ribosomes, and in turn access to actively translating mRNA, specifically within neurons of the mPFC-AcbSh pathway. RNA sequencing of the immunoprecipitated polysomes was performed to identify a circuit-based genetic profile that is associated with Susceptibility or Resistance to ABA. This type of approach represents a critical first step in the pursuit of understanding the *causes* of functional changes in reward and cognitive processes involved in pathological weight loss. Our findings pave the way for validation and manipulation of specific genes in the ABA model in future studies to identify causal molecular mechanisms.

## Materials and Methods

### Animals and housing

All animals were obtained from the Monash Animal Research Platform (MARP; Clayton, VIC, Australia). Female Sprague-Dawley rats (*n*=28) were 7 weeks of age on arrival in the laboratory. Animals were group-housed and acclimated to the 12-hour light/dark cycle (lights off at 1100h) for 7 days in a temperature (22-24°C) and humidity (30-50%) controlled room before experiments commenced. A male rat was individually housed in all experimental rooms to facilitate synchronization of oestrous cycle of the female rats (Cora, Kooistra, & Travlos, 2015). All experimental procedures (see **Figure 1** for timeline) were conducted in accordance with the Australian Code for the care and use of animals for scientific purposes and approved by the Monash Animal Resource Platform Ethics Committee (ERM 29143).

**Figure 1.**
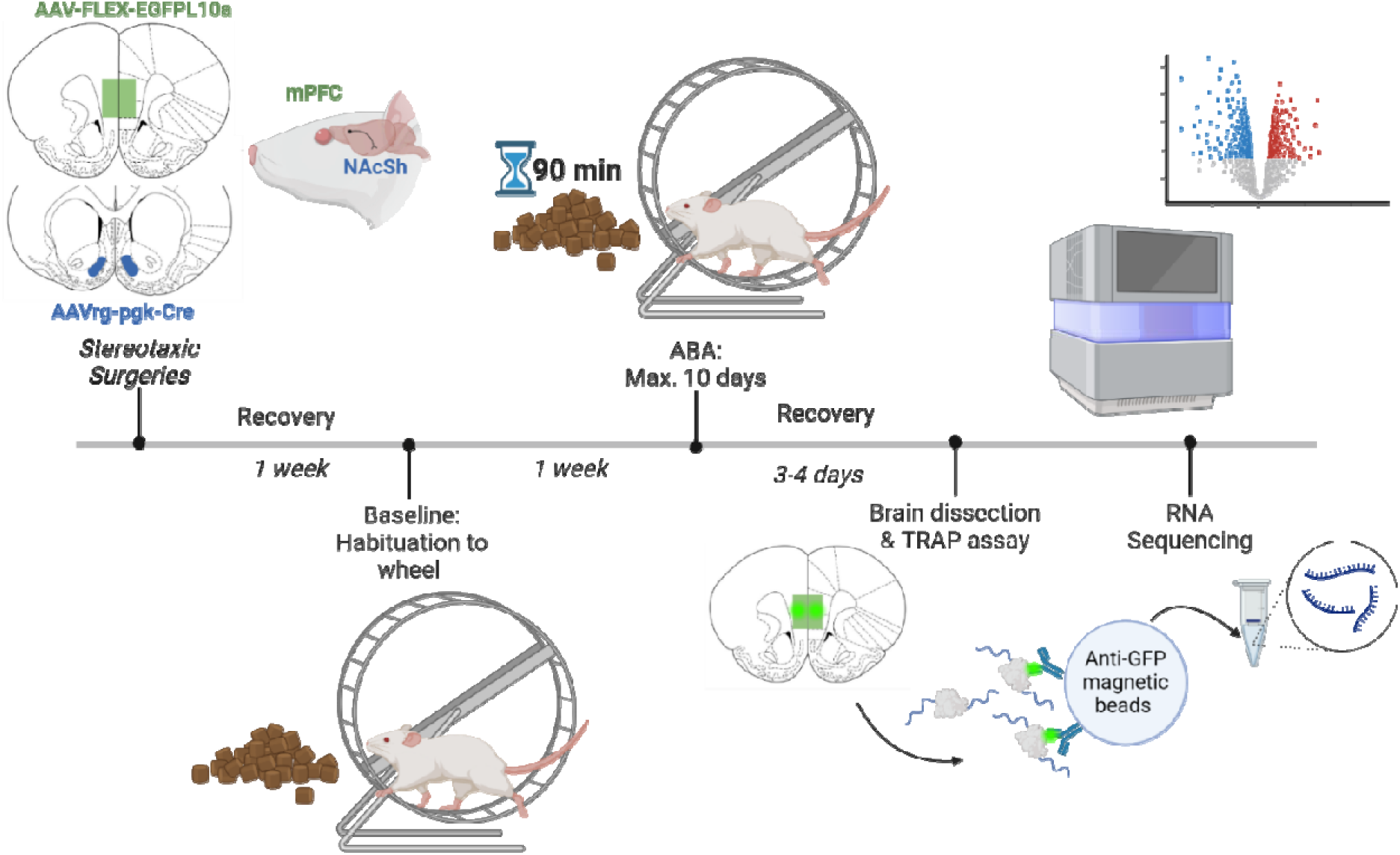
Timeline of experiments. Female Sprague-Dawley rats were injected with AAV-FLEX-EGFPL10a into medial prefrontal cortex (mPFC) and retrogradely transporting AAVrg-pgk-Cre into nucleus accumbens shell (NAcSh). Following one week of recovery, rats were habituated to running wheels with *ad libitum* food access for 7 days to determine baseline body weight and running wheel activity. Activity-based anorexia (ABA) conditions commenced with food access limited to 90 minutes per day and lasted for a maximum of 10 days or until rats reached <80% baseline body weight. Rats were allowed to recover to baseline body weight with *ad libitum* food access for 3 to 4 days before brain dissection. Tissue from mPFC was used in the TRAP assay to extract labelled RNA, followed by RNA sequencing and analyses.

### Stereotaxic surgeries

Stereotaxic surgeries were performed under anaesthesia (2.5-3% isoflurane in oxygen) using a stereotaxic apparatus (Kopf Instruments, CA, USA) and micropipettes pulled from borosilicate glass (0.64mm; Drummond Scientific, PA, USA) with a tip of ∼40µm. All animals were injected bilaterally with 250nL of retro-Cre (AAVrg-pgk-Cre; Addgene, #24593) into AcbSh (from bregma and brain surface: anteroposterior: +1.6mm, mediolateral: ± 0.7mm, dorsoventral: −6.6mm), followed by bilateral injection of 250nL of the Cre-dependent TRAP construct (AAV5-FLEX-EGFPL10a; Addgene #98747) into the mPFC (from bregma and brain surface: anteroposterior: +2.8mm, mediolateral: ± 0.5mm, dorsoventral: −3.2mm). Viral constructs were infused over 5 minutes (50nl/min), with pipette tips left at the coordinates for a further 5 minutes to allow virus dispersal. Meloxicam (2mg/kg; Boehringer Ingelheim, Germany) was subcutaneously injected into animals during surgery. Suture clips were used to close the surgery sites. Animals were group-housed with *ad libitum* food and water access post-surgery for 7 days to recover and allow for viral expression (see **Figure 2** for validation of expression and **Supplementary methods**).

**Figure 2.**
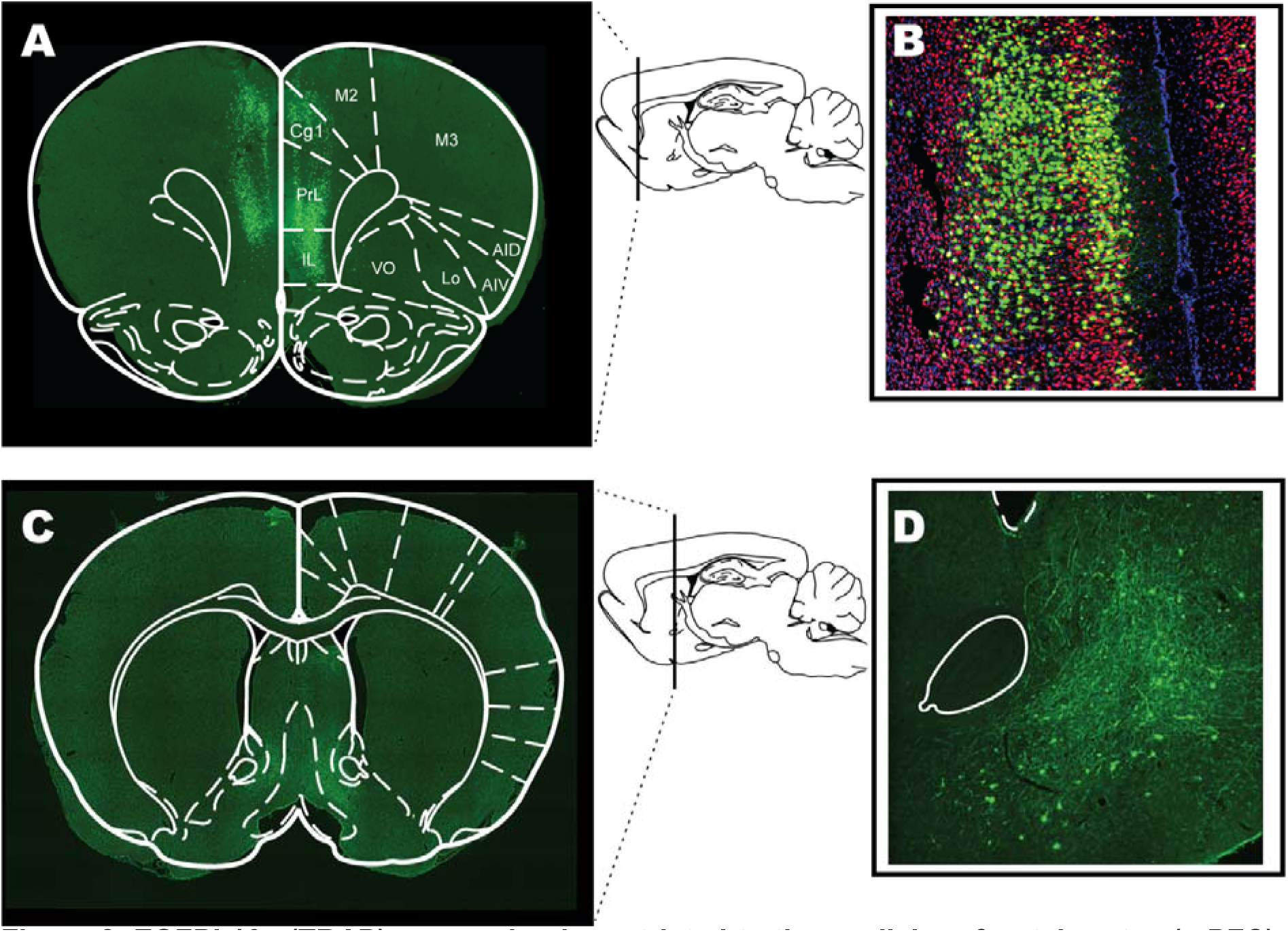
EGFPL10a (TRAP) expression is restricted to the medial prefrontal cortex (mPFC). (**A**) The mPFC region labelled with EFGP collected for RNA sequencing with atlas overlay (Bregma 3.20mm). (**B**) Micrograph (20X) of the mPFC midline with eGFPL10a (green), NeuN (red) and DAPI (blue) immunofluorescence. (**C**) The medial AcbSh region confirming terminal staining in the mPFC-AcbSh pathway with atlas overlay (Bregma 0.48mm). (**D**) Micrograph (20X) of terminal EGFP staining in the AcbSh [markup to orient to the anterior commissure bundle (solid line) and lateral ventricle (dashed line)].

### Activity-based anorexia (ABA)

Rats were individually housed in transparent living chambers with a removable food basket and a running wheel (Lafayette Instruments, IN, USA). Rats were allowed to habituate to the living chamber with *ad libitum* food access and locked running wheel for 1 day, followed by habituation to the unlocked running wheel for 7 days to determine baseline activity. Running wheel activity was recorded in revolutions by the Scurry Activity Wheel Software (Lafayette Instruments, IN, USA) with data extracted in 1h time bins. During ABA, food access was restricted to 90 minutes per days at the onset of dark phase (1100-1230h), with body weight measurements taken daily at 1100h. Running in the hour before the feeding window (1000-1100h) was considered as food anticipatory activity (FAA) (Milton et al., 2021; Milton et al., 2022). Time restricted food access persisted for a maximum of 10 days or until animals reached <80% of baseline body weight (ABA criterion), the baseline measure being that taken on Day 0 (prior to the first food restriction period). Following body weight recovery to at least 100% of baseline with *ad libitum* food access (minimum of 3 days to regain baseline weight), rats were euthanised with 300mg/kg sodium pentobarbitone (Lethabarb; Virbac, Australia), followed by rapid dissection of the mPFC region performed in dissection buffer containing 1x HBSS, 2.5mM HEPES-KOH (pH=7.4), 35mM glucose, 4mM NaHCO_3_ and 100 µg /ml cycloheximide (CHX). Dissected tissue was snap frozen in liquid nitrogen and stored at −80 °C until use.

### Translating Ribosome Affinity Purification (TRAP) assay and RNA Sequencing

TRAP was performed based on previously described protocols (Heiman et al., 2014; Ip et al., 2019; Nectow et al., 2017), followed by RNA extraction using the RNeasy Micro Kit (Qiagen, The Netherlands). Twelve samples were selected for sequencing according to weight loss profiles, in which equal numbers (*n*=4/group) were either clearly Resistant to developing ABA, rapidly lost weight under conditions of ABA and were removed from the experimental paradigm after 3-4 days (Susceptible) or exhibited an Intermediate phenotype, in which they lost weight to reach the removal criterion but more slowly than Susceptible rats (6-9 days). Total RNA samples underwent quality control by bioanalyser and the resulting library pool was QC’d by Qubit, Bioanalyzer and qPCR. Libraries were made by individual first strand synthesis to add the i7 index sequences indicated in **Supplementary Table 1**.

### Analysis

The raw sequencing reads were adapter trimmed using Trim Galore (v0.6.7; https://github.com/FelixKrueger/TrimGalore) with standard parameters. Reads were aligned to the Rat reference genome (Rattus_norvegicus.mRatBN7) with STAR (v 2.7.9a) (Dobin et al., 2013) and quantified using Salmon (v 1.9.0) (Patro, Duggal, Love, Irizarry, & Kingsford, 2017). The reference genome was downloaded from Ensembl, along with the gene annotation in a .gtf file. MultiQC (v 3.9.5) (Ewels, Magnusson, Lundin, & Käller, 2016) was used to generate the summary of the final quality metrics for all datasets. Differential expression analysis was carried out using edgeR (Robinson, McCarthy, & Smyth, 2010) along with the RUVseq (Risso, Ngai, Speed, & Dudoit, 2014) to remove unwanted noise in the data. Genes with 1 count per million in at least three samples per group were selected to perform the tests of differential expression. Significant differentially expressed genes (DEGs) were selected using log-fold-change of ±1 and FDR cut-off of 0.05 and were used for performing gene-set enrichment analysis (GSEA). The Rat Genome Database pathway and gene ontology (GO) analysis utilities were used (Petri et al., 2011). GSEA was used to identify classes of genes that are over-represented in a large set of genes or proteins, and may have an association with different phenotypes. GO analyses were summarized at the level of Biological process (BP), Cellular Component (CC), and Molecular Function (MF). Pathway analysis results were plotted using the online platform SRplot (http://www.bioinformatics.com.cn/SRplot) (Tang et al., 2023). Specific details of quality metrics from the MultiQC report are available on request.

### Statistical Analysis

Statistical analyses were performed using GraphPad Prism (GraphPad Software, San Diego, CA, USA). Statistical significance was considered as *p*<.05. Analyses including two-tailed unpaired t-test, one-way and two-way analysis of variance (ANOVA) with Tukey’s or Bonferroni’s post hoc multiple comparisons were used according to the number of groups in the ABA data. Volcano plots displaying DEGs and GO analyses for up and downregulated genes used a negative log FDR of >1.3 to determine statistical significance (*p<*.05).

## Results

### Susceptible and Resistant rats demonstrate distinct behavioural phenotypes in ABA

In order to confirm the TRAP expression within mPFC before the onset of ABA exposure, we used immunohistochemistry to validate that expression was restricted to neurons projecting to the AcbSh (**Figure 2**). In accordance with our previously characterized susceptibility criterion (C. J. Foldi et al., 2017; Milton et al., 2021; Milton et al., 2018), Resistant rats showed a slower body weight loss rate and maintained >80% of their baseline body weight for the entire 10-day ABA exposure period [**Figure 3A-D; B**, *p* = 0.0045; **C**, Experimental day *F*(10,74) = 15.0, *p* < 0.0001; ABA Susceptibility *F*(1,10) = 12.5, *p*= 0.0054; **D**, *p* = 0.0037]. Consistent with their ability to maintain body weight, Resistant rats consumed significantly more during the 90 minutes of food access throughout the 10-day period [**Figure 3E-F**; **E**, Experimental day *F*(9,63) = 9.46, *p* < 0.0001; ABA Susceptibility *F*(1,10) = 2.59, *p* = 0.1384; **F**, *p*= 0.0099], but did not demonstrate an increased level of food anticipatory activity (FAA; **Figure 3H**, *p*= .3029), contrary to previous reports (Milton et al., 2021). Resistant rats also maintained a relatively stable and moderate level of daily running wheel activity, whereas Susceptible rats showed a clear activity spike at the onset of food restriction [**Figure 3G**; ABA: Experimental day *F*(10,63) = 3.39, *p* = 0.0013; ABA Susceptibility *F*(1,10) = 10.9, *p*= 0.007].

**Figure 3.**
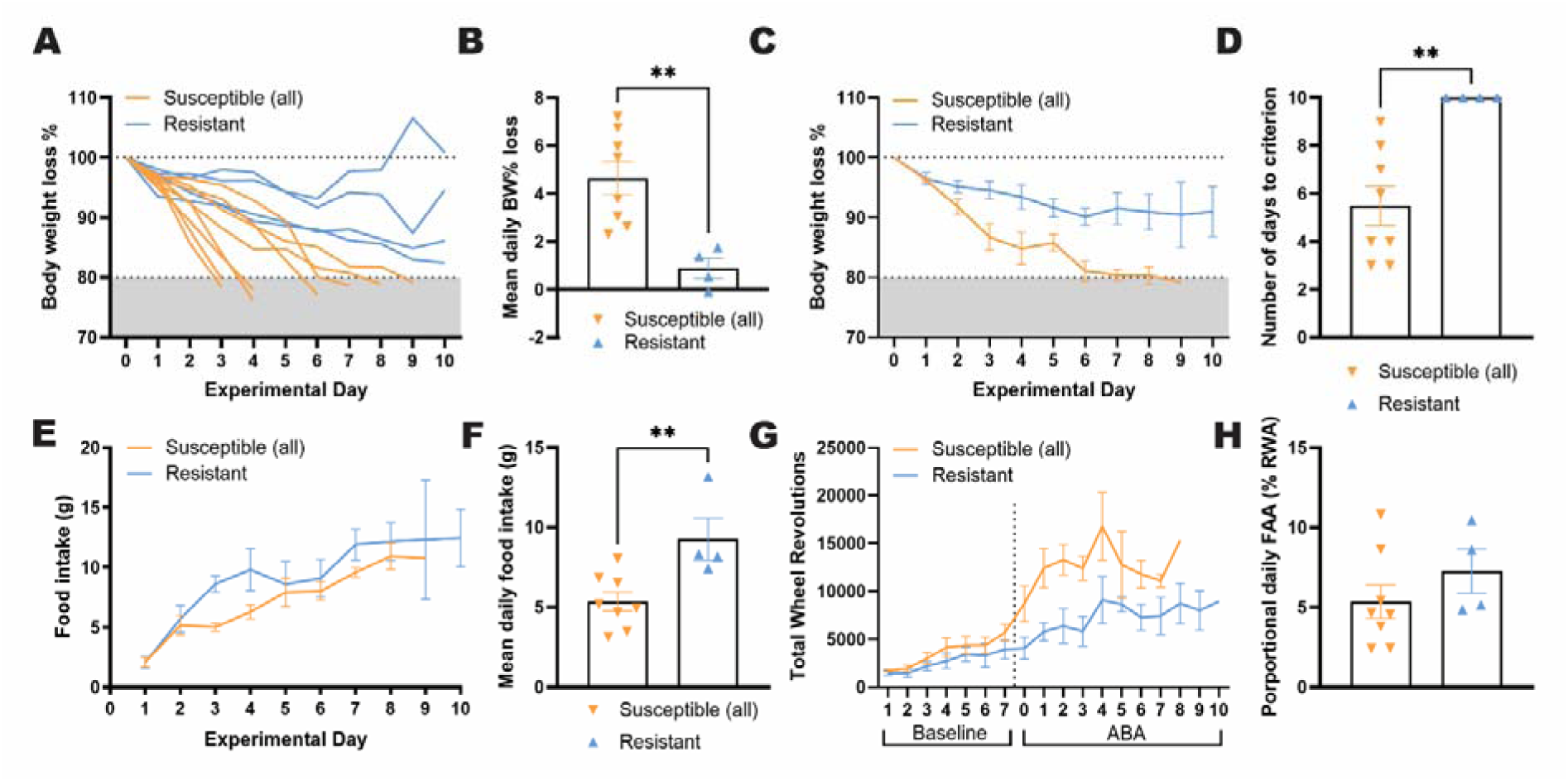
Primary ABA outcome measures for rats Susceptible and Resistant to weight loss. As expected based on susceptibility criteria, Resistant rats were able to maintain body weight above the threshold of 80% baseline body weight for the entire 10-day period, whereas Susceptible rats were not (**A**) and demonstrated a more rapid weight loss rate [**B-D**; **B**, *t*(10) = 3.640, *p* = 0.0045; **C**, Experimental day *F*(10,74) = 15.0, *p* < 0.0001; ABA Susceptibility *F*(1,10) = 12.5, *p*= 0.0054; **D**, *t*(10) = 3.770, *p* = 0.0037]. Resistant rats also ate more food within the time window than Susceptible rats during the 10-day ABA period [**E-F; E,** Experimental day *F*(9,63) = 9.46, *p* < 0.0001; ABA Susceptibility *F*(1,10) = 2.59, *p* = 0.1384; **F**, *t*(10) = 3.173, *p* = 0.0099]. Overall running wheel activity was significantly higher in Susceptible rats [**G;** Baseline: Experimental day *F*(6,60) = 9.01, *p* < 0.0001; ABA Susceptibility *F*(1,10) = 1.05, *p*= 0.3288; Interaction *F*(6,60) = 0.420, *p* = 0.8629; ABA: Experimental day *F*(10,63) = 3.39, *p* = 0.0013; ABA Susceptibility *F*(1,10) = 10.9, *p*= 0.0079], contributing to their improved body weight maintenance. However, motivated running in anticipation of food [**H**, *t*(10) = 1.086, *p* = 0.3029] was not significantly elevated in Resistant rats. Grouped data show mean ± SEM, with individual data point on the bar graphs. ** *p* < 0.01. ABA, Activity-based anorexia; BW, Body weight; FAA, Food anticipatory activity.

### Differentially expressed genes align with data from GWAS and implicate both metabolic and synaptic processes in pathological weight loss

Motivated by the complex roles of the cortico-striatal (mPFC-AcbSh) pathway during cognitive flexibility *versus* reinforcement learning in ABA rats, next we investigated whether stable molecular phenotypes in this pathway are associated with susceptibility to excessive exercise and weight loss. To examine enduring (and perhaps intrinsic) transcriptomic differences within the mPFC-AcbSh pathway between the two phenotypes, we harnessed the TRAP technology to selectively isolate ribosome-bound mRNA from this pathway after body weight recovery (at least to 100% of baseline). This approach was chosen to disambiguate compensatory mechanisms from possible circuit-specific differences underlying susceptibility to ABA. edgeR analysis between groups revealed 1424 differentially expressed genes (DEGs) between Susceptible and Resistant rats, highlighting important transcriptional differences associated with Resistance to ABA within this neural circuit (-log_10_FDR > 1.3, **Figure 4A**). To further understand the biological significance of DEGs, gene ontology (GO) analysis revealed that DEGs were significantly enriched in pathways related to oxidative phosphorylation and associated with neurodegenerative diseases, including Parkinson’s, Alzheimer’s and Huntington’s diseases (**Figure 4B)**. The major molecular and cellular-related themes that emerged were in genes associated with mitochondrial and cytoskeletal functions, cellular organisation and protein complex assembly (**Figure 4C-E)**. More specifically, among these genes we found that upregulated genes were mainly involved in metabolic processes (*Atp5mkI:* LogFC= 1.50*; Uqcr10*: LogFC= 1.32) and downregulated genes were involved in cytoskeletal, postsynaptic and axonal functions (blue box; **Figure 4A**). Interestingly, 5 of the DEGs were previously identified in GWAS as AN-associated genes (Watson et al., 2019). Of these, *Zc3h10*, *Myl6*, *Ikzf4* and *Erlec1* were upregulated in Resistant compared to Susceptible ABA rats, whereas *Smarcc2* was downregulated in the Resistant group. These genes have functions in thermoregulation (Yi et al., 2019), inflammatory responses (Liu et al., 2023) and endoplasmic reticulum stress (Misiewicz et al., 2013).

**Figure 4:**
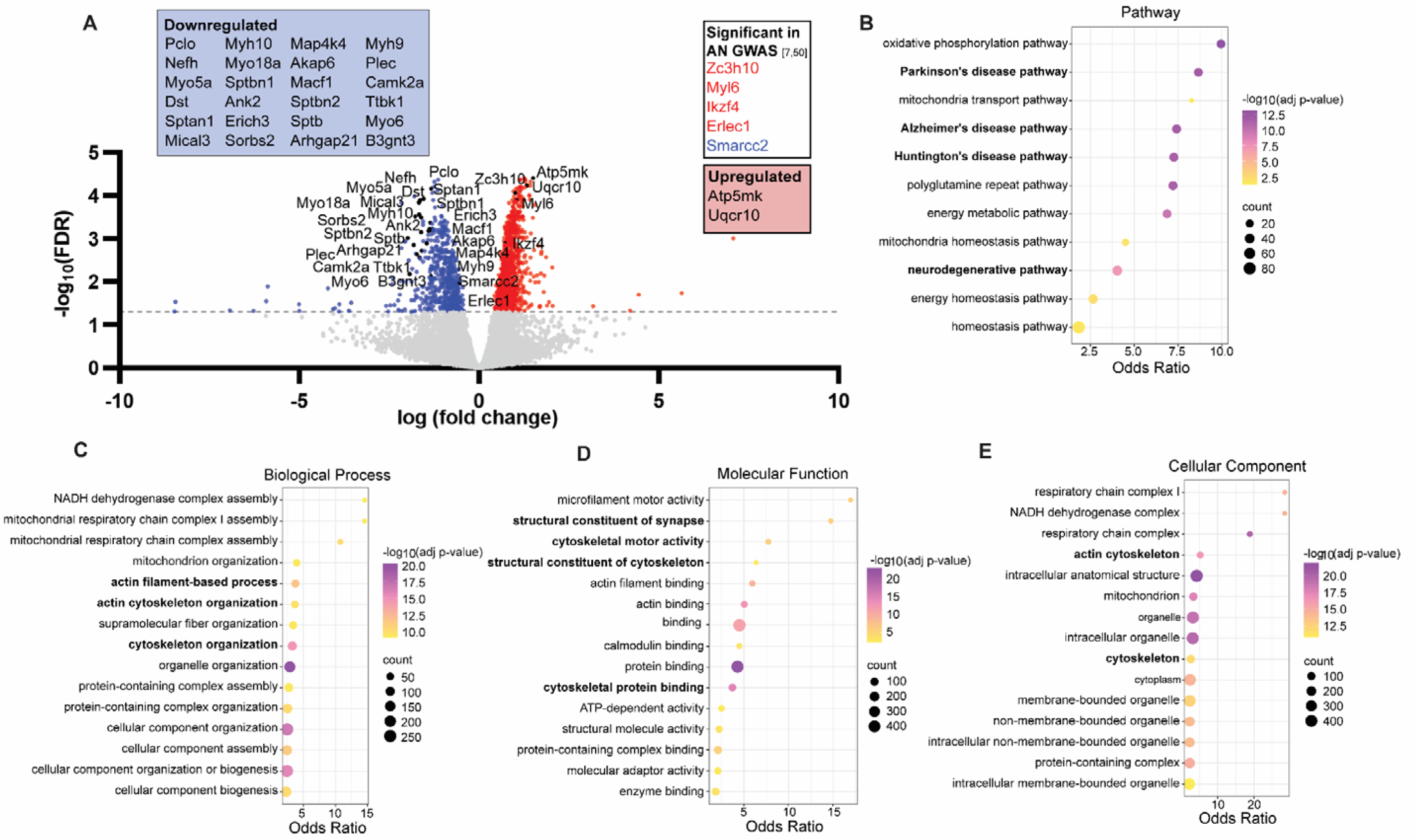
Rats that are Resistant to ABA show distinct gene expression profiles in striatum projecting mPFC neurons compared to Susceptible rats. Volcano plot of 1424 significant differentially expressed genes (DEGs) between Susceptible and Resistant rats (**A**). Grey dashed line represents a −log_10_(FDR) = 1.3, the threshold for statistical significance (*p <*.05). The blue dots denote all significantly downregulated genes observed in the Resistant, the red dots denote all significantly upregulated genes in the Resistant group, and grey dots represent genes that differed in expression between groups that did not reach statistical significance. Gene names are indicated for the most significant and stable (i.e. the least between-sample variability) upregulated (2) or downregulated (24) genes and the 5 genes that are significant in human genome-wide associated studies for anorexia nervosa (Watson et al., 2019 & Duncan et al., 2017). Pathways that were significantly (corrected *p*-value < 0.05) enriched in Susceptible vs. Resistant rats are involved in metabolic functions and neurodegenerative diseases (**B**). The top 15 biological process (**C**), molecular function (**D**), and cellular component (**E**) Gene Ontology (GO) terms enriched in DEGs predominantly relate to mitochondrial and cytoskeletal functions. The vertical axis of the bubble plots represents the pathway name or GO term, which is used to describe function of a gene. The horizontal axis represents the Odds Ratio, which is the likelihood of a pathway or GO term to be associated with a set of genes of interest compared to a reference set of genes. A higher odds ratio represents a stronger enrichment of the term or pathway. The colour scale for each bubble plot (**B-E**) indicates different thresholds of the *p*-value, whereby a lighter colour represents a greater significance. In addition, the bubbles size represents the number of genes observed to be differentially expressed within each pathway or term. Terms with a corrected *p*-value < 0.05 and an odds ratio between 1.3 and 40 were included.

### A subgroup of Susceptible rats shows an “Intermediate” behavioural phenotype

Through our extensive experience with the ABA model, we have noticed phenotypic differences within those rats Susceptible to ABA, particularly with respect to the *rate* of body weight loss. We were therefore interested to understand whether differences in weight loss trajectory were related to gene expression within this cortico-striatal pathway. Here, we distinguish Susceptible rats as those that lost weight rapidly (i.e., reached the threshold for ABA susceptibility criterion in <4 days) and Intermediate rats as those that still reached this criterion, albeit at a slower rate (**Figure 5A,-D; B**, Intermediate vs Susceptible *p* = 0.0003; Intermediate vs Resistant *p* = 0.0096; Susceptible vs Resistant *p* < 0.0001; **C**, *p* = 0.0079, **D**, Intermediate vs Susceptible *p* = 0.0002; Intermediate vs Resistant *p* = 0.0049; Susceptible vs Resistant *p* < 0.0001). As described above, Resistant rats ate significantly more when it was available than Susceptible rats [**Figure 5E-F**; **E,** Experimental day *F*(9,63) = 9.06, *p* < 0.0001; ABA Susceptibility *F*(2,9) = 1.22, *p* = 0.3389; **F**, *p* = 0.0084], and Intermediate rats consumed food in amounts between the two other groups. Interestingly, Resistant and Intermediate rats engaged in the same level of food anticipatory activity, and although this was reduced for Susceptible rats, the difference was not significant (**Figure 5H**, Intermediate vs Susceptible *p* = 0.1550; Resistant vs Susceptible *p* =0.1430). Examination of the profiles of daily running wheel activity showed a clear spike in activity at the onset of food restriction (dashed line) for Susceptible rats, whereas Intermediate rats exhibit a more gradual increase in wheel running and Resistant rats maintain a relatively constant rate of exercise [**Figure 5G** ABA: Experimental day *F*(10,63) = 3.58, *p* = 0.0008; ABA Susceptibility *F*(2,9) = 8.24, *p*= 0.0092].

**Figure 5:**
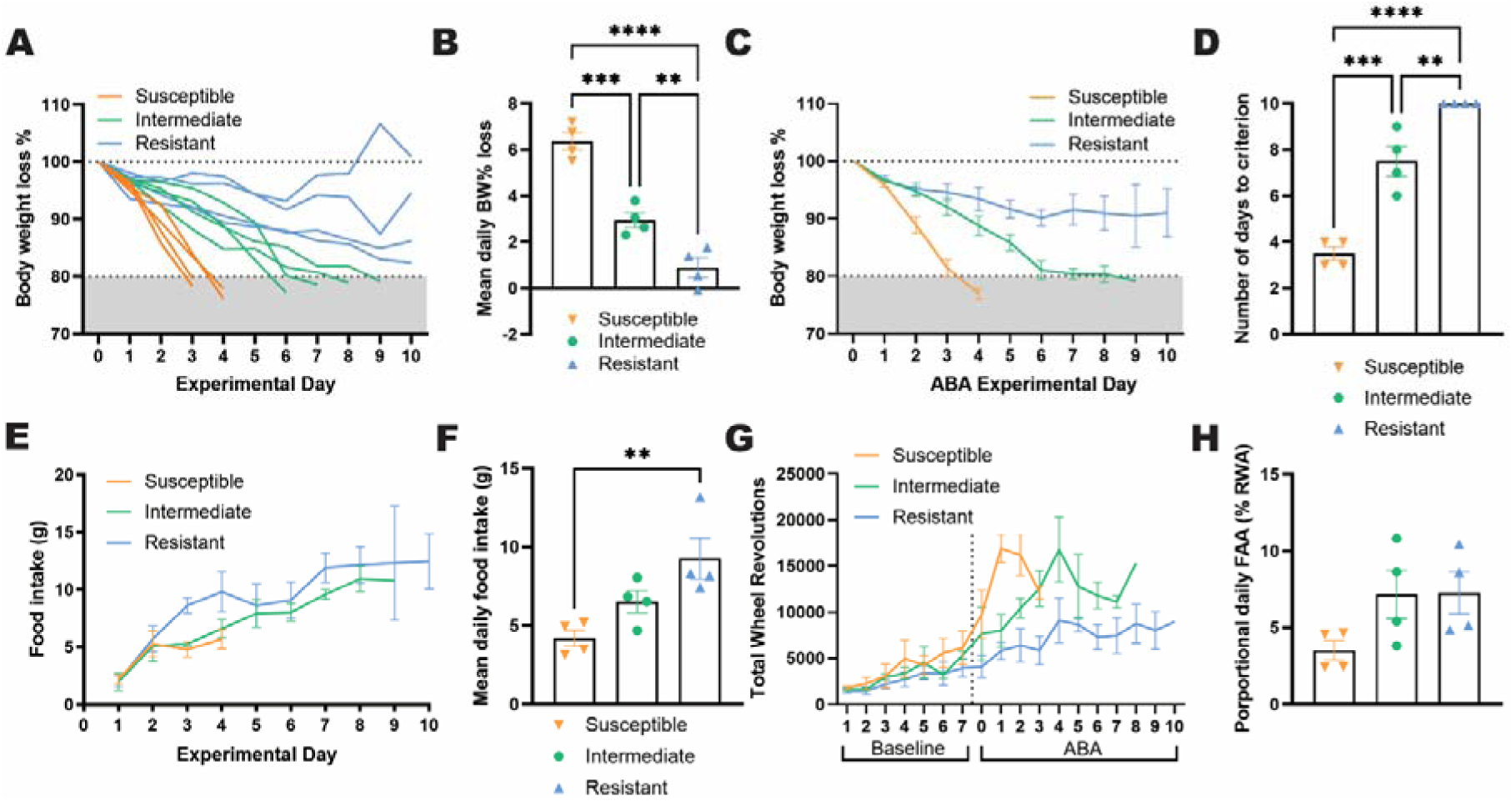
Separation of ABA rats based on trajectory of weight loss reveals an intermediate phenotype. While Susceptible rats lost weight rapidly, Intermediate rats reached the threshold for ABA susceptibility criterion, albeit at a slower rate [**A-D**, **B**, *F*(2,9) = 54.3, *p* < 0.0001, Intermediate vs Susceptible *p* = 0.0003; Intermediate vs Resistant *p* = 0.0096; Susceptible vs Resistant *p* < 0.0001; **C**, Experimental day *F*(10,74) =15.3, p < 0.0001, ABA Susceptibility *F*(2,9) = 8.68, *p* = 0.0079; **D**, *F*(2,9) = 64.5, *p* < 0.0001, Intermediate vs Susceptible *p* = 0.0002; Intermediate vs Resistant *p* = 0.0049; Susceptible vs Resistant *p* < 0.0001] and engaged in feeding at levels between the two other groups [but not significantly different to either; **E-F**;**E,** Experimental day *F*(9,63) = 9.06, *p* < 0.0001; ABA Susceptibility *F*(2,9) = 1.22, *p* = 0.3389 ;**F** *F*(2,9) = 7.84, *p* = 0.0107, Resistant > Intermediate *p* = 0.1364, Intermediate > Susceptible *p* = 0.219]. Examination of the profiles of daily running wheel activity showed a clear spike in activity at the onset of food restriction (dashed line) for Susceptible rats, whereas Intermediate rats exhibit a more gradual increase in wheel running [**G,** Baseline: Experimental day *F*(6, 54) = 11.50, *p* < 0.0001; ABA Susceptibility *F*(2, 9) = 0.7325 *p*= 0.5073; Interaction *F*(12,54) = 0.7031, *p* = 0.7415; ABA: Experimental day *F*(10,63) = 3.58, *p* = 0.0008; ABA Susceptibility *F*(2,9) = 8.24, *p*= 0.0092]. Interestingly, Resistant and Intermediate rats engaged in the same level of motivated running in anticipation of food (FAA) [**H**, *F*(2,9) = 2.889, *p* = 0.1074, Intermediate vs Susceptible *p* = 0.1550, Resistant vs Intermediate *p* = 0.9984]. Grouped data show mean ± SEM, with individual data point on the bar graphs. ** *p* < 0.01, *** *p* < 0.001, \*\*\*\**p* < 0.0001. ABA, Activity-based anorexia; BW, Body weight; FAA, Food anticipatory activity.

With this weight loss separation in mind, we examined the differences between two subgroups of Susceptible rats compared to Resistant group and identified genes that were differentially expressed between groups as well as those that overlapped. Overall, 651 DEGs were identified as overlapping and substantially fewer DEGs were identified between Resistant and Intermediate subgroups than those identified between Resistant and Susceptible, supporting their separation into discrete categories (**Figure 6A**). Within these DEGs, 170 genes exclusively differed in the Intermediate group, whereas 471 exclusively differed in the Susceptible group, further supporting the subgroup delineation. Moreover, GO analysis revealed zero overlap between the two comparator groups in terms of the top 15 biological processes, with Resistant and Susceptible rats mostly differing in genes related to metabolic and mitochondrial functions, whereas Resistant and Intermediate rats were most different by structural constituent and cytoskeletal functions. Specifically, the Intermediate group had upregulated genes associated with neurodevelopmental disorders (e.g. *Shank1-3*) whereas the Susceptible group were more genetically distinct to resistant rats, with upregulation of genes related to synaptic function (e.g. *Camk2a*) and downregulation of genes involved in satiety (e.g. *Cck*), inflammatory processes (e.g. *Ccl27*), and neuronal growth (e.g. *Bdnf*).

**Figure 6:**
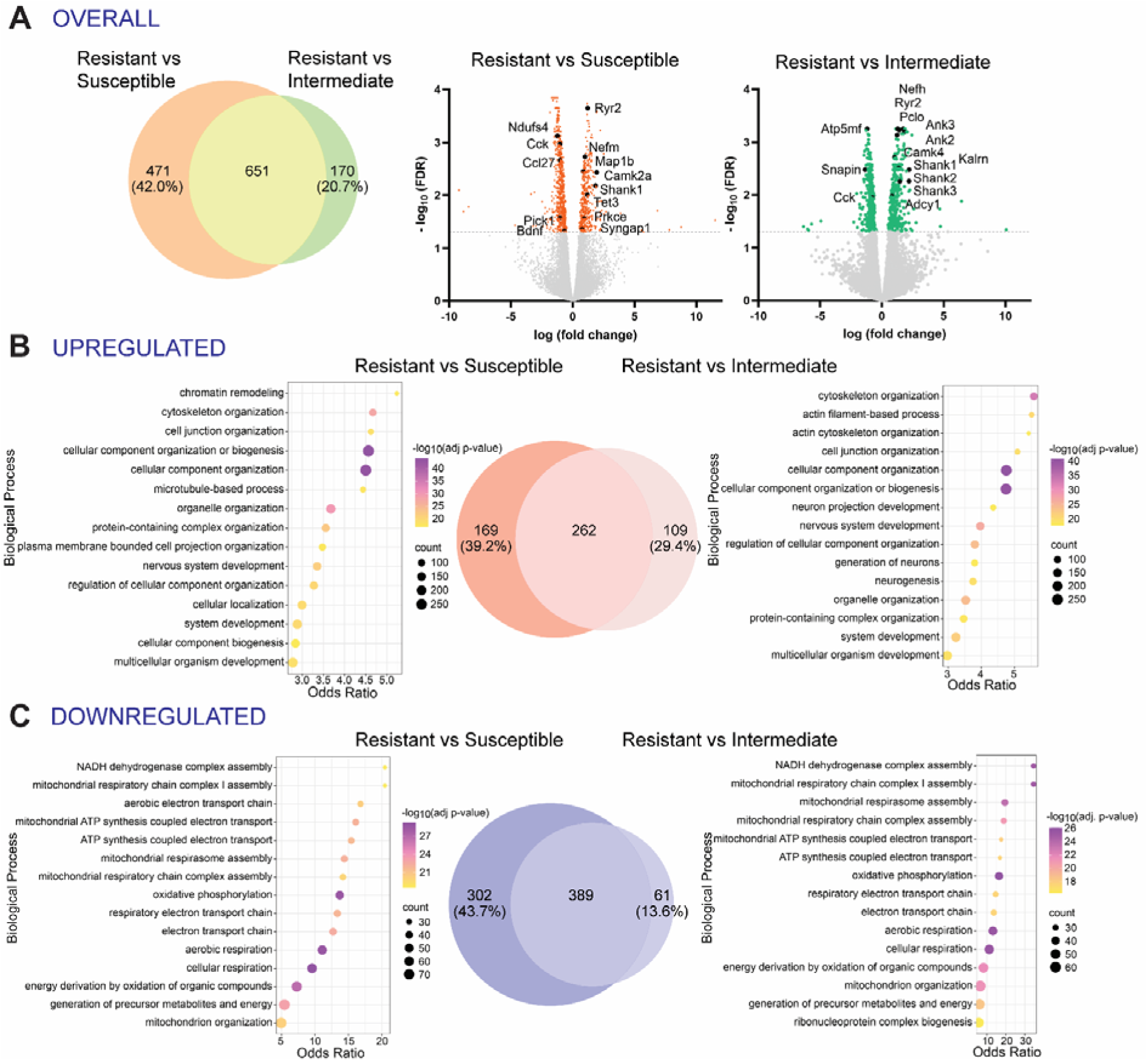
Gene expression profiles for ABA Resistant rats, compared to Susceptible or Intermediate subgroups. Venn diagram shows overlap of differentially expressed genes (DEGs) in Susceptible and/or Intermediate compared to Resistant rats (**A**), noting a large group of genes (651) are common to both comparisons and that there are fewer DEGs exclusively found in the Resistant vs. Intermediate comparison (170) than the Resistant vs Susceptible comparison (471). Volcano plots of DEGs for Resistant compared to Susceptible or Intermediate subgroup. The significant DEGs [-log_10_(FDR) > 1.3] are indicated in orange or green, with gene names of interest indicated for genes that are related to neurodevelopmental disorders, synaptic function, satiety, inflammatory processes and feeding behaviours, all processes relevant to the development of ABA in rats and AN in humans. Venn diagram shows overlapping of DEGs that are upregulated in Susceptible (left) or Intermediate (right) rats compared to Resistant rats (**B**), with 39.2% of upregulated DEGs that are exclusive to Susceptible but not Intermediate. 10 of the Top 15 biological process terms enriched in the upregulated DEGs common between the two comparisons (i.e. Susceptible vs Resistant and Intermediate vs Resistant - see bubble plots). With respect to downregulated DEGs, the Venn diagram shows that 43.7% of identified genes are exclusive to the Susceptible vs Resistant group comparison and do not overlap with those in the Intermediate vs Resistant group comparison (**C**). Here, 14 of the Top 15 biological process terms enriched in the downregulated DEGs are common to both groups (see bubble plots), which are involved in metabolic and mitochondrial functions and suggests that these processes are a shared feature of both “weight losing” groups.

We were also interested in examining the up and downregulated DEGs between the Susceptible and Intermediate subgroups compared to the Resistant group separately. Similar to the overlap in DEGs, there was broad overlap between the Susceptible and Intermediate comparisons with Resistant rats, in which 262 upregulated genes overlapped (**Figure 6B**) and 389 downregulated genes overlapped (**Figure 6C**), with fewer DEGs identified between Resistant and Intermediate subgroups than those identified between Resistant and Susceptible. In addition, 10 of the top 15 biological processes enriched for upregulated genes in both comparisons (Susceptible vs Resistant and Intermediate vs Resistant) were the same, and 14 out of the top 15 biological processes enriched for downregulated genes were common to both Susceptible and Intermediate comparisons with Resistant rats.

### Susceptible and Intermediate subgroups show no significant differentially expressed genes

To determine whether Susceptible and Intermediate rats showed DEG within this same pathway, we directly compared the profile of these subgroups. Strikingly, no genes were differentially expressed using the statistically corrected value for significance, indicating the two subgroups of “ABA susceptible” rats do not display detectable differences in expression at the single gene level.

## Discussion

In this study, we used a novel viral-based TRAP technique to reveal transcriptional changes within the cortico-striatal neuronal circuit that are associated with susceptibility to pathological weight loss in ABA rats. These changes were associated with metabolic, mitochondrial and neuronal functions and independent of current calorie deficit, since assessment occurred after weight recovery. Upregulated genes in Resistant rats were enriched for mitochondrial function, in line with a recent report of mitochondrial damage leading to oxidative stress in ABA rats (Bhasin et al., 2023), while downregulated genes were involved in cytoskeletal, postsynaptic and axonal functions. Overall, these findings point to *decreased excitability of the mPFC-AcbSh circuits* as a molecular signature that may confer resistance to weight loss. This is consistent with our previous study showing that chemogenetic *suppression* of this pathway caused resistance to pathological weight loss during ABA (Milton et al., 2021) and the evidence of elevated activity of PFC regions in humans weight recovered from AN (Ehrlich et al., 2015).

GWAS provide an excellent starting point for understanding the genetic architecture of complex psychiatric disorders. Our study builds upon GWAS findings by investigating how gene expression (i.e. the “message”) is altered, rather than only the DNA (i.e. the “code”). One risk gene identified in the largest GWAS investigation in AN cases (Watson et al., 2019), *Erlec1*, was significantly downregulated in Susceptible compared to Resistant rats, but this was not significant when comparing Intermediate to Resistant rats, suggesting that differences observed in weight loss trajectories might involve dysregulated protein metabolism. Unfolded or misfolded proteins in the endoplasmic reticulum (ER) are recognised by lectins including *Erlec1,* and an impairment in this recognition process could disrupt ER homeostasis and cause tissue inflammation and endocrine dysregulation (Lemmer, Willemsen, Hilal, & Bartelt, 2021). Notably, *Erlec1* demonstrates reliable patterns of under-expression in central nodes within fear and reward circuits, suggesting that genes linked with AN and vulnerability to rapid weight loss in ABA may confer risk by altering processes related to fear- and reward-learning (S. B. Murray et al., 2023), consistent with reports of persistent alterations in these processes after weight gain in AN (Stuart B. Murray et al., 2018) and with the role of this cortico-striatal circuit in punishment learning (conditioned suppression) (Piantadosi et al., 2020).

In addition, genes associated with feeding behaviour and reward processing, including *Bdnf*, were significantly downregulated in Susceptible rats compared to Resistant rats, but not for the Intermediate weight losing subgroup. *Bdnf* mRNA expression is reduced in mPFC by scheduled feeding in mice (Ho, Klenotich, McMurray, & Dulawa, 2016) and BDNF serum concentration is reduced in patients with AN (Nilsson et al., 2020). These data support the possibility that differences in weight loss trajectory in ABA could result from altered reward-related feeding behaviour (C. J. Foldi et al., 2017). Intriguingly, the *Cck* gene, encoding the peptide hormone cholecystokinin (CCK), was downregulated in both Susceptible and Intermediate rats, compared to Resistant rats. The direction of effect here seems counterintuitive when considering the role of CCK in signalling satiety (Moran, 2000), however it aligns with the observation of reduced circulating CCK in patients with AN (Baranowska, Radzikowska, Wasilewska-Dziubinska, Roguski, & Borowiec, 2000). Thus, considering gene expression reported here is restricted to the cortico-stratal circuitry, the downregulation we show may relate to the way that “higher order” cortical neurons that express CCK mRNA in the rat (Burgunder & Young, 1990) regulate appetite, through behavioural mechanisms such as attention and impulse control.

The view that separate genetic profiles are involved in the genesis of rapid versus slower weight loss in ABA rats is supported to some extent by the GSEA. Focusing on the top significantly enriched pathways associated with the Susceptible or Intermediate ABA phenotype (see **Supplementary Figure 1**), it is clear that dysregulation of genes associated with mitochondrial function (transport, dynamics and homeostasis) is common to both “weight losing” groups, whereas dysregulation of genes involved in neurodegenerative disease pathways was specific to rapid weight losers (i.e. Susceptible) and not significantly associated with the Intermediate phenotype. Individuals that respond to ABA conditions with rapid weight loss could be separated in future studies to investigate whether common neurochemical disturbances seen in Parkinson’s Disease (i.e. loss of dopamine neurons) or Alzheimer’s Disease (i.e. loss of acetylcholine neurons) play a role in the development or maintenance of AN-like behaviours. A better understanding of these synergies could lead to more effective treatments, in light of the fact that pharmacotherapies approved for neurodegenerative diseases, such as donepezil (Aricept), have been proposed as a feasible possibility for treating AN (Bucay, 2009), and recent advances in understanding the AN- relevant mechanisms in mouse models (Favier et al.). Interestingly, although the Enriched Pathways and Biological Processes were distinct when comparing Susceptible or Intermediate rats to Resistant rats, the Cellular Components and Molecular Functions identified as significantly dysregulated were largely overlapping (see **Supplementary Figures 2-3**). Consistent with the significant co-morbidity between AN and psychiatric disorders including depression, anxiety, substance use and obsessive-compulsive disorder (OCD) (Salbach-Andrae et al., 2008), genetic correlations between these have also been revealed via single-nucleotide polymorphism (SNP)- based methods. In particular, AN has the strongest (positive) genetic correlation with OCD (Watson et al., 2019) and shares similar alterations in PFC expression (Song et al., 2021), suggesting a common dysregulated functional pathway. Direct evidence of genome-wide significant risk that is shared between AN and OCD has not been identified, however, several genes have been suggested to drive the association, including *Lrrc16a* and *Kit* (Yilmaz et al., 2020), both involved in cell signalling pathways within the gastrointestinal tract. Moreover, 109 pleiotropic loci were identified in a cross-disorder GWAS of psychiatric disorders (including AN and OCD), in which the *Dcc* gene, the product of which plays a key role in guiding axonal growth during development, was significantly associated with all eight psychiatric disorders (Lee et al., 2019). In addition, this study identified common genetic associations between AN and major psychiatric disorders including schizophrenia, bipolar disorder and major depression, as well as psychological traits including mood instability, neuroticism and intelligence, of which several (*Csnk2b, Ctsa, Gpx1, P4htm and Tcf7l2)* were differentially expressed by the Susceptible and Resistant rats, suggesting that the ABA model might capture genetic underpinnings of other psychiatric traits that predispose individuals to AN. Despite the fact that the most translatable feature of the ABA model is the development of compulsive exercise, which is an intermediate phenotype of shared genetic risk for both OCD and AN (Yilmaz et al., 2023), the putative genes that link AN to OCD were not found to be differentially dysregulated in the present study. This could be due to the fact that specific risk loci shared by AN and OCD have yet to be pinpointed (Bulik et al., 2022) or that excessive exercise is driven not by altered mPFC-AcbSh pathway activity but by disrupted function in other cortico-striatal pathways, for example, those terminating in the dorsal striatum (Conn, Huang, Gorrell, & Foldi, 2024).

## Limitations and future directions

While our study provides novel insight into transcriptomic signatures associated with vulnerability to ABA, several limitations constrain the interpretation and immediate translational impact of our findings. Chief among these is that the current work stops short of establishing causality between differentially expressed genes (DEGs) and AN-like phenotypes. Although our results identify promising molecular targets within PFC–AcbSh projecting neurons, future studies should employ gene manipulation strategies (e.g., viral overexpression or knockdown) to determine whether altering expression of candidate DEGs can modulate ABA susceptibility. Specifically, manipulating expression in vulnerable rats to promote resilience (or vice versa) would allow for a more definitive demonstration of causal relevance. Additionally, pharmacological validation of identified pathways, if feasible, could provide important preclinical support for novel therapeutic strategies. Despite this, our results support the use of the ABA model in investigating causal genetic factors to AN and provide directions for the development of translationally relevant genetic models of AN. Because of the complex nature of psychiatric genetics, an appropriate genetic model of AN is unlikely to involve a single gene manipulation, thus, developing novel genetic models that incorporate ABA with multiple specific and inducible manipulations of AN-associated genes, or genetic variants such as copy number variants (CNVs; (Walker et al., 2024)) or short structural variants (SSVs; (Berthold et al., 2022)) could provide a better opportunity for understanding the causal relationship between genetic factors, altered cognitive functions and feeding or exercise pathologies.

A related limitation is the absence of molecular validation for key DEGs. As with all unbiased RNA sequencing techniques, our approach carries an inherent risk of false positives and negatives. Although we applied appropriate statistical correction and prioritised biologically plausible targets for discussion, confirmatory experiments are warranted. In particular, in situ hybridization or immunohistochemistry in conjunction with the retrograde labelling approach used here would help establish spatial and cell-type specificity of gene expression changes within PFC–NAc projection neurons. With respect to behavioural controls, we clarify that our study design explicitly sought to contrast rats that were susceptible versus resistant to ABA. That is, within the same experimental conditions of food restriction and running wheel access, the tendency to lose weight differentiates vulnerable rats from those who maintain weight. In this context, susceptible animals serve as the relevant behavioural control and demonstrate the same trajectory of weight loss and excessive exercise as we have reported numerous times (Huang et al., 2023; Milton et al., 2021; Milton et al., 2018; Milton et al., 2022). Nonetheless, comparing gene expression profiles with a separate food-restricted, non-ABA group would provide important information to disentangle the contribution of weight loss *per se* from the motivational or behavioural drivers of the ABA phenotype.

In conclusion, this study suggests that weight loss in ABA results from hyperexcitability in this cortico-striatal pathway, which is normalised in rats resistant to weight loss. The increased expression of axonal and synaptic genes could indicate that too much neuronal activity in this circuit may cause maladaptive behaviour. Future studies should aim to validate this interpretation with circuit-based studies that include markers of pre- and post-synaptic densities, in which the hypothesis would be that a greater “synaptic load” would be colocalised with the mPFC-AcbSh projecting cells in ABA Susceptible compared to Resistant rats.

## Acknowledgements

We acknowledge the technical assistance with quality control and novel RNA sequencing pipelines developed and performed by Dr Trevor Wilson, Monash Health Translational Precinct, Medical Genomics Facility. We acknowledge the Monash Microimaging Platform for infrastructure used to obtain immunofluorescence micrographs. We also acknowledge Biorender.com for elements of the figure schematics.

## Data Availability Statement

Data are available from the corresponding author upon reasonable request.

## Notes

**Funding:** This work was supported by a National Health and Medical Research Council (NHMRC) of Australia Ideas Grant (GNT2001722) awarded to CJF. K.H. is supported by a Monash Graduate Scholarship (MGS) and K.C. is supported by a Monash Faculty Early Career Postdoctoral Fellowship.

**Conflict of interest:** The authors declare no conflict of interest.

### Competing Interest Statement

The authors have declared no competing interest.

### Summary of Updates

Modified some of the text to make clear this is a first step in the pursuit of uncovering a genetic signature of susceptibility to pathological weight loss. Removed one figure and moved one figure from the supplement to the main text.

## References

Baranowska, B., Radzikowska, M., Wasilewska-Dziubinska, E., Roguski, K., & Borowiec, M. (2000). Disturbed release of gastrointestinal peptides in anorexia nervosa and in obesity. *Diabetes*, Obesity and Metabolism, 2(2), 99–103. 10.1046/j.1463-1326.2000.00070.x

Barbarich-Marsteller, N. C., Underwood, M. D., Foltin, R. W., Myers, M. M., Walsh, B. T., Barrett, J. S., & Marsteller, D. A. (2013). Identifying novel phenotypes of vulnerability and resistance to activity-based anorexia in adolescent female rats. Int J Eat Disord, 46(7), 737–746. doi:10.1002/eat.22149

Beeler, J. A., & Burghardt, N. S. (2021). Activity-based Anorexia for Modeling Vulnerability and Resilience in Mice. Bio Protoc, 11(9), e4009. doi:10.21769/BioProtoc.4009

Berthold, N., Pytte, J., Bulik, C. M., Tschochner, M., Medland, S. E., & Akkari, P. A. (2022). Bridging the gap: Short structural variants in the genetics of anorexia nervosa. Int J Eat Disord, 55(6), 747–753. doi:10.1002/eat.23716

Bhasin, H., O’Brien, S. C., Cordner, Z. A., Aston, S. A., Tamashiro, K. L. K., & Moran, T. H. (2023). Activity-based anorexia in adolescent female rats causes changes in brain mitochondrial dynamics. Physiology & Behavior, 261, 114072. 10.1016/j.physbeh.2022.114072

Bucay, A. H. (2009). Donepezil (aricept) as a treatment for anorexia nervosa: a very feasible therapeutic possibility. Expert Opinion on Investigational Drugs, 18(5), 569–571. doi:10.1517/13543780902810360

Bulik, C. M., Coleman, J. R. I., Hardaway, J. A., Breithaupt, L., Watson, H. J., Bryant, C. D., & Breen, G. (2022). Genetics and neurobiology of eating disorders. Nat Neurosci, 25(5), 543–554. doi:10.1038/s41593-022-01071-z

Burgunder, J. M., & Young, W. S., 3rd. (1990). Cortical neurons expressing the cholecystokinin gene in the rat: distribution in the adult brain, ontogeny, and some of their projections. J Comp Neurol, 300(1), 26–46. doi:10.1002/cne.903000104

Carrera, O., Gutiérrez, E., & Boakes, R. A. (2006). Early handling reduces vulnerability of rats to activity-based anorexia. Dev Psychobiol, 48(7), 520–527. doi:10.1002/dev.20175

Cha, J., Ide, J. S., Bowman, F. D., Simpson, H. B., Posner, J., & Steinglass, J. E. (2016). Abnormal reward circuitry in anorexia nervosa: A longitudinal, multimodal MRI study. Human Brain Mapping, 37(11), 3835–3846. 10.1002/hbm.23279

Chesney, E., Goodwin, G. M., & Fazel, S. (2014). Risks of all-cause and suicide mortality in mental disorders: a meta-review. World Psychiatry, 13(2), 153–160. doi:10.1002/wps.20128

Conn, K., Huang, K., Gorrell, S., & Foldi, C. J. (2024). A transdiagnostic and translational framework for delineating the neuronal mechanisms of compulsive exercise in anorexia nervosa. International Journal of Eating Disorders, 57(7), 1406–1417. 10.1002/eat.24130

Cora, M. C., Kooistra, L., & Travlos, G. (2015). Vaginal Cytology of the Laboratory Rat and Mouse: Review and Criteria for the Staging of the Estrous Cycle Using Stained Vaginal Smears. Toxicol Pathol, 43(6), 776–793. doi:10.1177/0192623315570339

Cox, J., & Witten, I. B. (2019). Striatal circuits for reward learning and decision-making. Nature Reviews Neuroscience, 20(8), 482–494. doi:10.1038/s41583-019-0189-2

Dobin, A., Davis, C. A., Schlesinger, F., Drenkow, J., Zaleski, C., Jha, S., … Gingeras, T. R. (2013). STAR: ultrafast universal RNA-seq aligner. Bioinformatics, 29(1), 15–21. doi:10.1093/bioinformatics/bts635

Ehrlich, S., Geisler, D., Ritschel, F., King, J. A., Seidel, M., Boehm, I., … Kroemer, N. B. (2015). Elevated cognitive control over reward processing in recovered female patients with anorexia nervosa. J Psychiatry Neurosci, 40(5), 307–315. doi:10.1503/jpn.140249

Ewels, P., Magnusson, M., Lundin, S., & Käller, M. (2016). MultiQC: summarize analysis results for multiple tools and samples in a single report. Bioinformatics, 32(19), 3047–3048. doi:10.1093/bioinformatics/btw354

Favier, M., Janickova, H., Justo, D., Kljakic, O., Runtz, L., Natsheh, J. Y., … El Mestikawy, S. Cholinergic dysfunction in the dorsal striatum promotes habit formation and maladaptive eating. (1558-8238 (Electronic)).

Fladung, A. K., Schulze, U. M., Schöll, F., Bauer, K., & Grön, G. (2013). Role of the ventral striatum in developing anorexia nervosa. Transl Psychiatry, 3(10), e315. doi:10.1038/tp.2013.88

Foldi, C. J. (2024). Taking better advantage of the activity-based anorexia model. Trends in Molecular Medicine, 30(4), 330–338. 10.1016/j.molmed.2023.11.011

Foldi, C. J., Milton, L. K., & Oldfield, B. J. (2017). The Role of Mesolimbic Reward Neurocircuitry in Prevention and Rescue of the Activity-Based Anorexia (ABA) Phenotype in Rats. Neuropsychopharmacology, 42(12), 2292–2300. doi:10.1038/npp.2017.63

Hancock, S., & Grant, V. (2009). Early maternal separation increases symptoms of activity-based anorexia in male and female rats. J Exp Psychol Anim Behav Process, 35(3), 394–406. doi:10.1037/a0014736

Heiman, M., Kulicke, R., Fenster, R. J., Greengard, P., & Heintz, N. (2014). Cell type–specific mRNA purification by translating ribosome affinity purification (TRAP). Nature Protocols, 9(6), 1282–1291. doi:10.1038/nprot.2014.085

Ho, E. V., Klenotich, S. J., McMurray, M. S., & Dulawa, S. C. (2016). Activity-Based Anorexia Alters the Expression of BDNF Transcripts in the Mesocorticolimbic Reward Circuit. PLoS One, 11(11), e0166756. doi:10.1371/journal.pone.0166756

Huang, K., Milton, L. K., Dempsey, H., Power, S. J., Conn, K. A., Andrews, Z. B., & Foldi, C. J. (2023). Rapid, automated, and experimenter-free touchscreen testing reveals reciprocal interactions between cognitive flexibility and activity-based anorexia in female rats. Elife, 12. doi:10.7554/eLife.84961

Ip, C. K., Zhang, L., Farzi, A., Qi, Y., Clarke, I., Reed, F., … Herzog, H. (2019). Amygdala NPY Circuits Promote the Development of Accelerated Obesity under Chronic Stress Conditions. Cell Metabolism, 30(1), 111–128.e116. 10.1016/j.cmet.2019.04.001

Joshi, A., Schott, M., la Fleur, S. E., & Barrot, M. (2022). Role of the striatal dopamine, GABA and opioid systems in mediating feeding and fat intake. Neuroscience & Biobehavioral Reviews, 139, 104726. 10.1016/j.neubiorev.2022.104726

Kawa, A. B., Hashimoto, J. G., Beutler, M. M., Guizzetti, M., & Wolf, M. E. (2025). Changes in nucleus accumbens core translatome accompanying incubation of cocaine craving. Neuropsychopharmacology. doi:10.1038/s41386-025-02112-4

Lee, P. H., Anttila, V., Won, H., Feng, Y.-C. A., Rosenthal, J., Zhu, Z., … Smoller, J. W. (2019). Genomic Relationships, Novel Loci, and Pleiotropic Mechanisms across Eight Psychiatric Disorders. Cell, 179(7), 1469–1482.e1411. doi:10.1016/j.cell.2019.11.020

Lemmer, I. L., Willemsen, N., Hilal, N., & Bartelt, A. (2021). A guide to understanding endoplasmic reticulum stress in metabolic disorders. Molecular Metabolism, 47, 101169. 10.1016/j.molmet.2021.101169

Liu, W., Wang, Z., Liu, S., Zhang, X., Cao, X., & Jiang, M. (2023). RNF138 inhibits late inflammatory gene transcription through degradation of SMARCC1 of the SWI/SNF complex. Cell Reports, 42(2). doi:10.1016/j.celrep.2023.112097

Milton, L. K., Mirabella, P. N., Greaves, E., Spanswick, D. C., van den Buuse, M., Oldfield, B. J., & Foldi, C. J. (2021). Suppression of Corticostriatal Circuit Activity Improves Cognitive Flexibility and Prevents Body Weight Loss in Activity-Based Anorexia in Rats. Biol Psychiatry, 90(12), 819–828. doi:10.1016/j.biopsych.2020.06.022

Milton, L. K., Oldfield, B. J., & Foldi, C. J. (2018). Evaluating anhedonia in the activity-based anorexia (ABA) rat model. Physiol Behav, 194, 324–332. doi:10.1016/j.physbeh.2018.06.023

Milton, L. K., Patton, T., O’Keeffe, M., Oldfield, B. J., & Foldi, C. J. (2022). In pursuit of biomarkers for predicting susceptibility to activity-based anorexia in adolescent female rats. Int J Eat Disord, 55(5), 664–677. doi:10.1002/eat.23705

Misiewicz, M., Déry, M.-A., Foveau, B., Jodoin, J., Ruths, D., & LeBlanc, A. C. (2013). Identification of a Novel Endoplasmic Reticulum Stress Response Element Regulated by XBP1 *. Journal of Biological Chemistry, 288(28), 20378–20391. doi:10.1074/jbc.M113.457242

Moran, T. H. (2000). Cholecystokinin and satiety: current perspectives. Nutrition, 16(10), 858–865. 10.1016/S0899-9007(00)00419-6

Murray, S. B., Rokicki, J., Sartorius, A. M., Winterton, A., Andreassen, O. A., Westlye, L. T., … Quintana, D. S. (2023). Brain-based gene expression of putative risk genes for anorexia nervosa. Mol Psychiatry, 28(6), 2612–2619. doi:10.1038/s41380-023-02110-2

Murray, S. B., Strober, M., Craske, M. G., Griffiths, S., Levinson, C. A., & Strigo, I. A. (2018). Fear as a translational mechanism in the psychopathology of anorexia nervosa. Neuroscience & Biobehavioral Reviews, 95, 383–395. 10.1016/j.neubiorev.2018.10.013

Nectow, A. R., Moya, M. V., Ekstrand, M. I., Mousa, A., McGuire, K. L., Sferrazza, C. E., … Schmidt, E. F. (2017). Rapid Molecular Profiling of Defined Cell Types Using Viral TRAP. Cell Rep, 19(3), 655–667. doi:10.1016/j.celrep.2017.03.048

Nilsson, I. A. K., Millischer, V., Göteson, A., Hübel, C., Thornton, L. M., Bulik, C. M., … Landén, M. (2020). Aberrant inflammatory profile in acute but not recovered anorexia nervosa. Brain, Behavior, and Immunity, 88, 718–724. 10.1016/j.bbi.2020.05.024

Patro, R., Duggal, G., Love, M. I., Irizarry, R. A., & Kingsford, C. (2017). Salmon provides fast and bias-aware quantification of transcript expression. Nat Methods, 14(4), 417–419. doi:10.1038/nmeth.4197

Petri, V., Shimoyama, M., Hayman, G. T., Smith, J. R., Tutaj, M., de Pons, J., … Team, R. G. D. (2011). The Rat Genome Database Pathway Portal. Database, 2011, bar010. doi:10.1093/database/bar010

Piantadosi, P. T., Yeates, D. C. M., & Floresco, S. B. (2020). Prefrontal cortical and nucleus accumbens contributions to discriminative conditioned suppression of reward-seeking. Learn Mem, 27(10), 429–440. doi:10.1101/lm.051912.120

Risso, D., Ngai, J., Speed, T. P., & Dudoit, S. (2014). Normalization of RNA-seq data using factor analysis of control genes or samples. Nat Biotechnol, 32(9), 896–902. doi:10.1038/nbt.2931

Robinson, M. D., McCarthy, D. J., & Smyth, G. K. (2010). edgeR: a Bioconductor package for differential expression analysis of digital gene expression data. Bioinformatics, 26(1), 139–140. doi:10.1093/bioinformatics/btp616

Salbach-Andrae, H., Lenz, K., Simmendinger, N., Klinkowski, N., Lehmkuhl, U., & Pfeiffer, E. (2008). Psychiatric comorbidities among female adolescents with anorexia nervosa. Child Psychiatry Hum Dev, 39(3), 261–272. doi:10.1007/s10578-007-0086-1

Scharner, S., & Stengel, A. (2021). Animal Models for Anorexia Nervosa—A Systematic Review. Frontiers in Human Neuroscience, 14. Retrieved from https://www.frontiersin.org/articles/10.3389/fnhum.2020.596381

Song, W., Wang, W., Yu, S., & Lin, G. N. (2021). Dissection of the Genetic Association between Anorexia Nervosa and Obsessive-Compulsive Disorder at the Network and Cellular Levels. Genes (Basel), 12(4). doi:10.3390/genes12040491

Tang, D., Chen, M., Huang, X., Zhang, G., Zeng, L., Zhang, G., … Wang, Y. (2023). SRplot: A free online platform for data visualization and graphing. PLoS One, 18(11), e0294236. doi:10.1371/journal.pone.0294236

Walker, A., Karlsson, R., Szatkiewicz, J. P., Thornton, L. M., Yilmaz, Z., Leppä, V. M., … Wray, N. R. (2024). Genome-wide copy number variation association study in anorexia nervosa. Molecular Psychiatry. doi:10.1038/s41380-024-02811-2

Watson, H. J., & Bulik, C. M. (2013). Update on the treatment of anorexia nervosa: review of clinical trials, practice guidelines and emerging interventions. Psychol Med, 43(12), 2477–2500. doi:10.1017/s0033291712002620

Watson, H. J., Yilmaz, Z., Thornton, L. M., Hübel, C., Coleman, J. R. I., Gaspar, H. A., … Bulik, C. M. (2019). Genome-wide association study identifies eight risk loci and implicates metabo-psychiatric origins for anorexia nervosa. Nat Genet, 51(8), 1207–1214. doi:10.1038/s41588-019-0439-2

Yi, D., Dempersmier, J. M., Nguyen, H. P., Viscarra, J. A., Dinh, J., Tabuchi, C., … Sul, H. S. (2019). Zc3h10 Acts as a Transcription Factor and Is Phosphorylated to Activate the Thermogenic Program. Cell Rep, 29(9), 2621–2633.e2624. doi:10.1016/j.celrep.2019.10.099

Yilmaz, Z., Halvorsen, M., Bryois, J., Yu, D., Thornton, L. M., Zerwas, S., … Crowley, J. J. (2020). Examination of the shared genetic basis of anorexia nervosa and obsessive-compulsive disorder. Mol Psychiatry, 25(9), 2036–2046. doi:10.1038/s41380-018-0115-4

Yilmaz, Z., Schaumberg, K., Halvorsen, M., Goodman, E. L., Brosof, L. C., Crowley, J. J., … Zerwas, S. C. (2023). Predicting eating disorder and anxiety symptoms using disorder-specific and transdiagnostic polygenic scores for anorexia nervosa and obsessive-compulsive disorder. Psychol Med, 53(7), 3021–3035. doi:10.1017/s0033291721005079

