## Supplementary for "Toward a genetic signature of resistance to activity-based anorexia in striatal projecting cortical neurons"

**Supplementary Table 1. Quality report for selected samples.** Note that while some samples (in bold) have lower than normal RIN values, the way in which the reads were collected indicate that this was caused by fragmentation (from bead-based physical immunoprecipitation) rather than by degradation of the samples (Trevor Wilson, *personal communication*).

| Sample number | Days to <80% BW criterion | Group | RNA Integrity Number (RIN) | BA conc [ng/µl] | i7 Index | Reads (M) |
|---|---|---|---|---|---|---|
| 1 | / | Resistant | 6.5 | 1.68 | TAAGGCGA | 34.1 |
| 3 | / | Resistant | 7.2 | 2.18 | TAAGGCGA | 88.5 |
| 4 | 4 | Susceptible | 7.4 | 1.59 | AGGCAGAA | 29.0 |
| 8 | 4 | Susceptible | 6.2 | 3.04 | TCCTGAGC | 20.8 |
| 9 | 6 | Intermediate | 7.7 | 10.75 | GGACTCCT | 27.0 |
| 11 | / | Resistant | 6.5 | 3.02 | TAGGCATG | 21.6 |
| 12 | 7 | Intermediate | 9.6 | 3.74 | CTCTCTAC | 38.8 |
| 15 | 3 | Susceptible | **3.4** | 1.81 | CGAGGCTG | 29.6 |
| 16 | 3 | Susceptible | **3.0** | 1.60 | AAGAGGCA | 109.9 |
| 23 | 8 | Intermediate | **3.6** | 5.97 | GTAGAGGA | 25.2 |
| 26 | 9 | Intermediate | **3.8** | 3.46 | GCTCATGA | 56.0 |
| 27 | / | Resistant | **4.8** | 0.51 | ATCTCAGG | 42.3 |

**Supplementary Methods:**

Immunohistochemical validation of TRAP viral expression

To validate the expression of the EGFPL10a (TRAP) construct, stereotaxic injections were performed as described. Two weeks later, rats (n=2) were euthanized with sodium pentobarbitone (Lethabarb 150mg/kg; Virbac, AU) and transcardially perfused with 200mL 0.9% saline followed by 200mL 4% paraformaldehyde in phosphate buffer. Brains were excised and postfixed in 4% paraformaldehyde in phosphate buffer overnight at 4°C, followed by submersion in 30% sucrose in phosphate buffer for 4 days. Brains were then sectioned at 35µm using a cryostat (CM1860; Leica Biosystems) and collected in a 1:4 series. A series of sections were immunolabeled with chicken anti-GFP (1:1000, Abcam) and guinea pig anti-NeuN (1:1000, Synaptic Systems) primary antibodies overnight at room temperature, followed by 90-minute incubation in Alexa-fluor conjugated secondary antibodies (1:1000, Alexa-fluor 488, Jackson ImmunoResearch; 1:1000, Alexa-fluor 594, Abcam) and 5-minute incubation in DAPI (1:5000, Sigma-Aldrich). Sections were imaged on a widefield microscope (Thunder Imager Live Cell & 3D Assay, Leica Microsystems, Germany).

Translating Ribosome Affinity Purification (TRAP) assay

To prepare the tissue for extraction of EGFP-tagged polysomes, the tissue was homogenised with a hand pestle mixer in lysis buffer [10 mM HEPES-KOH, 150 mM KCl, 5 mM MgCl2, 0.5 mM DTT, 100 µg/ml CHX, RNase inhibitor (RNasin, Promega, 40u/µl) and protease inhibitors (Roche, 1 tablet/ml)], followed by centrifugation at 2000g at 4°C for 10 minutes. The supernatant was then incubated with NP-40 working solution and DHPC on ice for 5 minutes, followed by centrifugation at

13,000 g for 10 minutes at 4°C. Following centrifugation, polysomes were immunoprecipitated from the supernatant by overnight incubation at 4°C with anti-GFP antibody-labelled Dynabeads (Invitrogen, California, USA). The bound RNA was collected with a magnetic rack and washed with a KCl wash buffer (10 mM HEPES-KOH, 350 mM KCl, 5 mM MgCl2, 1% NP-40, 0.5 mM DTT, 100 µg/ml CHX), followed by RNA extraction using the RNeasy Micro Kit (Qiagen, The Netherlands).

RNA Sequencing

Total RNA samples underwent quality control by bioanalyser and the resulting library pool was QC'd by Qubit, Bioanalyzer and qPCR. It had a mean size of 361bp at a concentration of 9.9nM. Sequencing was performed using NextSeq2000 on a P2 100cycle kit (cDNA reads generated 101nt), with read coverage up to 400M reads per run. A novel RNA-Seq pipeline (Version 01/09/2021) was used on the final library pool, loaded at 1000pM (based upon size adjusted qPCR) for on-board denaturation and clustering (Illumina Protocol; 1000000109376 v3 Nov2020) which resulted in a loading metric of 98.42 and 88.95% >Q30. 541.2 reads passed filter and 464 Million reads were assigned to indexes. Briefly, index is added during initial pA priming and pooled samples amplified using template switching oligo. P5 is added by tagmentation by Nextera transposase and PCR. On-board denaturing and clustering were conducted using 1000pM of library pool. Parsing and base calling was conducted (Dragen BCLConvert 3.7.4) and showed that Sample 22-2449 (Resistant) had the lowest amount of sample with 6.5ng total RNA. Since all samples are pooled for the multiplex method, this amount was used as starting amount for all samples.

**Supplementary Information: Definition of Gene Ontology terms**

Gene ontology (GO) describes gene functions in three aspects: Molecular Function, Cellular Component, and Biological Process. Molecular function describes the activity of the gene product at molecular level. Cellular component describes the location where the activity occurs in the cell. Biological process represents biological events contributed by molecular processes.

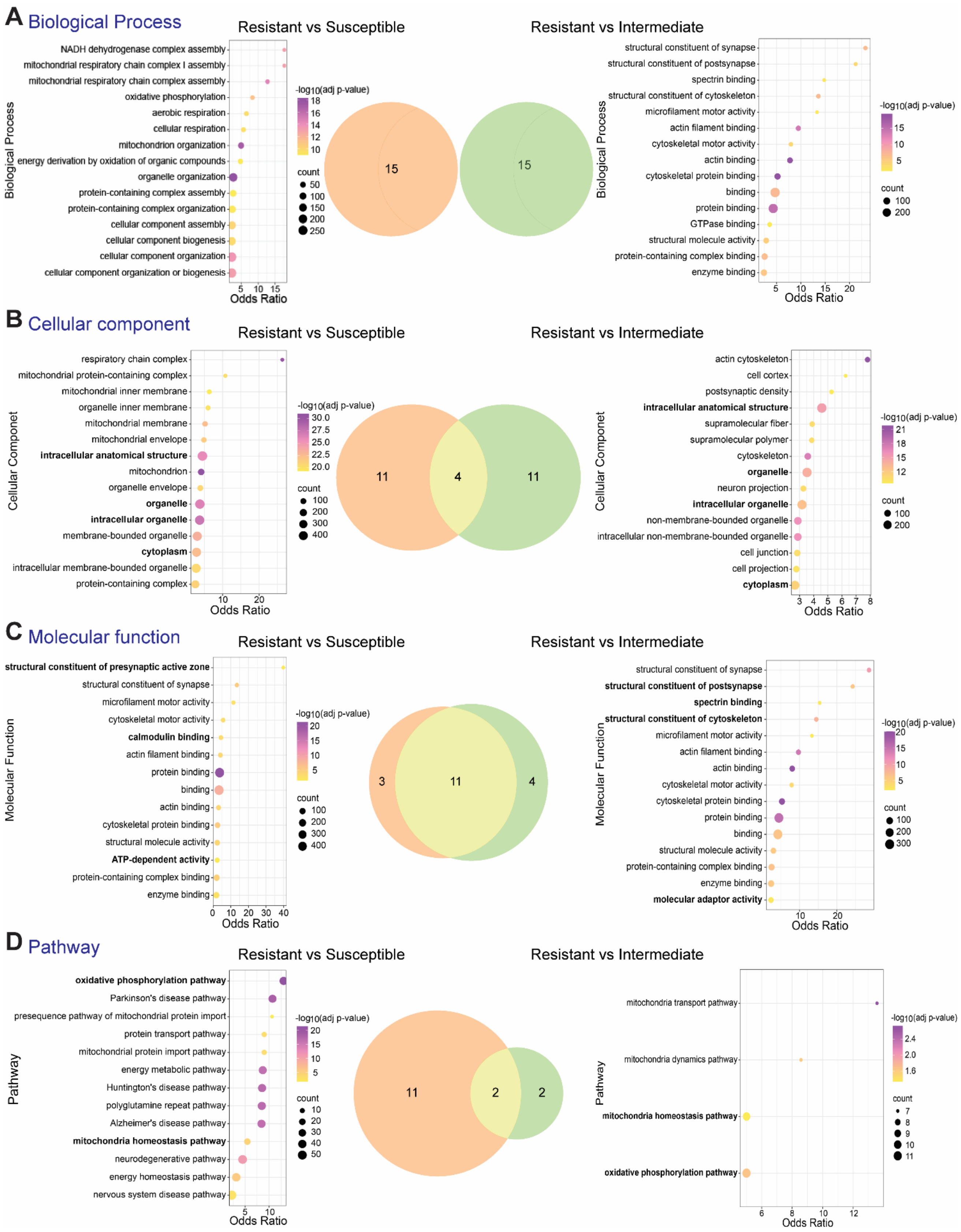
A Biological Process
Resistant vs Susceptible
Resistant vs Intermediate
NADH dehydrogenase complex assembly
mitochondrial respiratory chain complex I assembly
mitochondrial respiratory chain complex assembly
oxidative phosphorylation
aerobic respiration
cellular respiration
mitochondrion organization
energy derivation by oxidation of organic compounds
organelle organization
protein-containing complex assembly
protein-containing complex organization
cellular component assembly
cellular component biogenesis
cellular component organization
cellular component organization or biogenesis
Biological Process
Odds Ratio
-log10(adj p-value)
count
15
15
structural constituent of synapse
structural constituent of postsynapse
spectrin binding
structural constituent of cytoskeleton
microfilament motor activity
actin filament binding
cytoskeletal motor activity
actin binding
cytoskeletal protein binding
binding
protein binding
GTPase binding
structural molecule activity
protein-containing complex binding
enzyme binding
B Cellular component
Resistant vs Susceptible
Resistant vs Intermediate
respiratory chain complex
mitochondrial protein-containing complex
mitochondrial inner membrane
organelle inner membrane
mitochondrial membrane
mitochondrial envelope
intracellular anatomical structure
mitochondrion
organelle envelope
organelle
intracellular organelle
membrane-bounded organelle
cytoplasm
intracellular membrane-bounded organelle
protein-containing complex
Cellular Componet
11
4
11
actin cytoskeleton
cell cortex
postsynaptic density
intracellular anatomical structure
supramolecular fiber
supramolecular polymer
cytoskeleton
organelle
neuron projection
intracellular organelle
non-membrane-bounded organelle
intracellular non-membrane-bounded organelle
cell junction
cell projection
cytoplasm
C Molecular function
Resistant vs Susceptible
Resistant vs Intermediate
structural constituent of presynaptic active zone
structural constituent of synapse
microfilament motor activity
cytoskeletal motor activity
calmodulin binding
actin filament binding
protein binding
binding
actin binding
cytoskeletal protein binding
structural molecule activity
ATP-dependent activity
protein-containing complex binding
enzyme binding
Molecular Function
3
11
4
structural constituent of synapse
structural constituent of postsynapse
spectrin binding
structural constituent of cytoskeleton
microfilament motor activity
actin filament binding
actin binding
cytoskeletal motor activity
cytoskeletal protein binding
protein binding
binding
structural molecule activity
protein-containing complex binding
enzyme binding
molecular adaptor activity
D Pathway
Resistant vs Susceptible
Resistant vs Intermediate
oxidative phosphorylation pathway
Parkinson's disease pathway
presequence pathway of mitochondrial protein import
protein transport pathway
mitochondrial protein import pathway
energy metabolic pathway
Huntington's disease pathway
polyglutamine repeat pathway
Alzheimer's disease pathway
mitochondria homeostasis pathway
neurodegenerative pathway
energy homeostasis pathway
nervous system disease pathway
Pathway
11
2
2
mitochondria transport pathway
mitochondria dynamics pathway
mitochondria homeostasis pathway
oxidative phosphorylation pathway

**Supplementary Figure 1**. **Gene ontology terms and pathways enriched in differentially expressed genes for Resistant compared to Susceptible or Intermediate subgroups.** The top 15 significantly (corrected *p*-value < 0.05) enriched biological process (**A**), cellular components (**B**), molecular functions (**C**) gene ontology (GO) terms and pathways (**D**) are shown in bubble plots, with Venn diagrams depicting the specific overlap of these GO terms or pathways that were associated with the Susceptible (orange) or Intermediate (green) subgroups compared to Resistant rats. (**A**) The top 15 significant biological process enriched were entirely distinct for the comparisons between Susceptible versus Resistant and Intermediate versus Resistant groups. Differentially expressed genes (DEGs) in the Susceptible subgroup were most significantly associated with mitochondrial function and energy metabolism, whereas DEGs in Intermediate subgroup were mainly associated with cytoskeletal structure and protein interactions. (**B**) Cellular component terms included intracellular anatomical structure, organelle, intracellular organelle and cytoplasm that were common to both subgroups (see bold text term names), whereas mitochondrial structure and function are significantly dysregulated in Susceptible subgroup and cytoskeletal and neuronal structures are associated with the Intermediate subgroup. (**C**) The top 15 molecular function terms largely overlap between the two subgroups (11/15; cytoskeletal dynamics, synaptic function and protein binding), however, pre- and postsynaptic function differed between the comparison groups, whereby presynaptic function was exclusively associated with the Susceptible subgroup and postsynaptic function exclusively within the Intermediate subgroup (see bold terms). (**D**) Pathways associated mitochondrial function (transport, dynamics and homeostasis) are common to both Susceptible and Intermediate subgroups, whereas pathways involved in neurodegenerative diseases (Parkinson’s, Huntington’s and Alzheimer’s Disease) were only seen when comparing the Susceptible subgroup to Resistant rats.

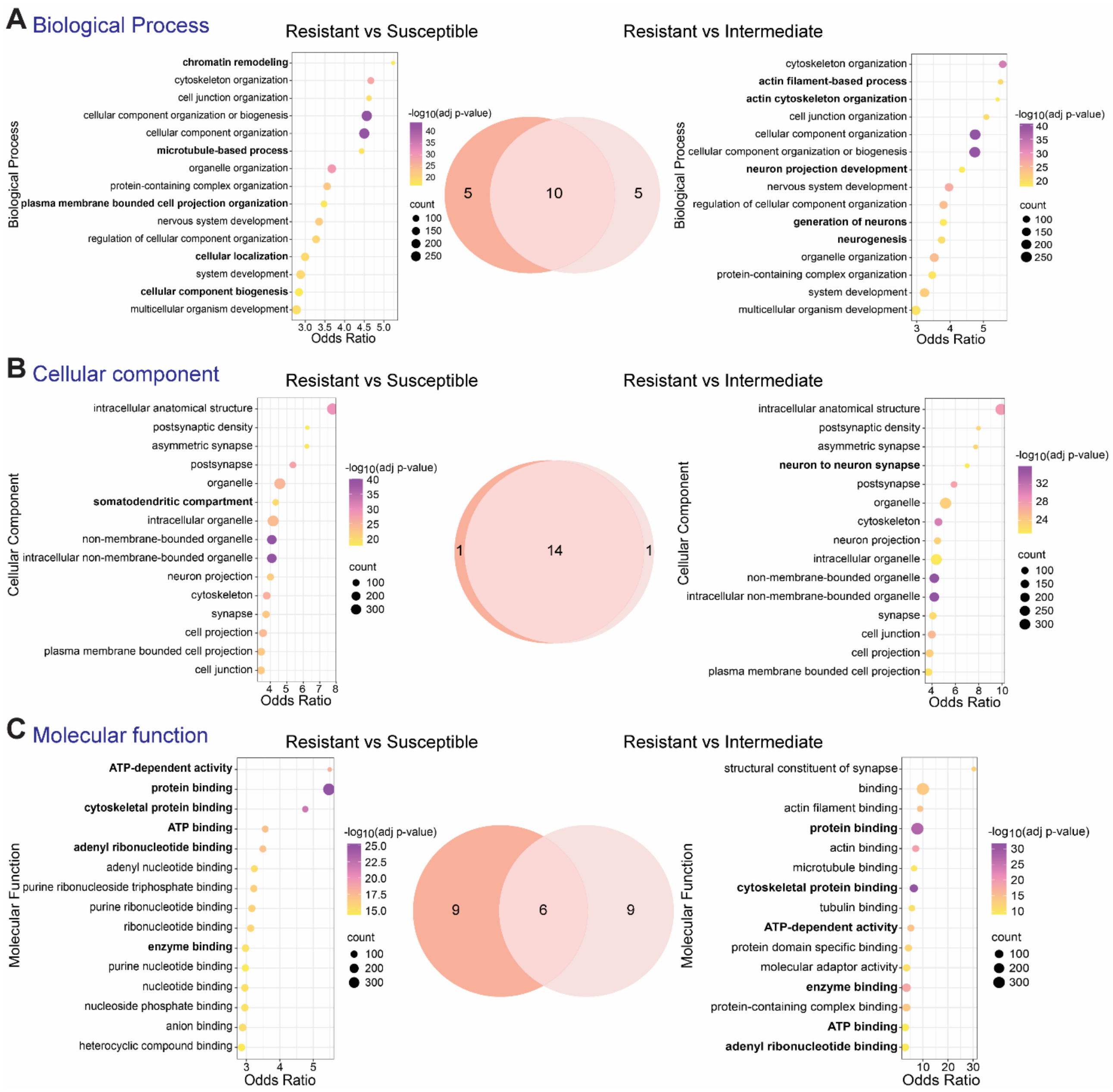
A Biological Process
Resistant vs Susceptible
Resistant vs Intermediate
Biological Process
chromatin remodeling
cytoskeleton organization
cell junction organization
cellular component organization or biogenesis
cellular component organization
microtubule-based process
organelle organization
protein-containing complex organization
plasma membrane bounded cell projection organization
nervous system development
regulation of cellular component organization
cellular localization
system development
cellular component biogenesis
multicellular organism development
3.0 3.5 4.0 4.5 5.0
Odds Ratio
-log10(adj p-value)
40 35 30 25 20
count
100 150 200 250
5 10 5
cytoskeleton organization
actin filament-based process
actin cytoskeleton organization
cell junction organization
cellular component organization
cellular component organization or biogenesis
neuron projection development
nervous system development
regulation of cellular component organization
generation of neurons
neurogenesis
organelle organization
protein-containing complex organization
system development
multicellular organism development
3 4 5
Odds Ratio
-log10(adj p-value)
40 35 30 25 20
count
100 150 200 250
B Cellular component
Resistant vs Susceptible
Resistant vs Intermediate
Cellular Component
intracellular anatomical structure
postsynaptic density
asymmetric synapse
postsynapse
organelle
somatodendritic compartment
intracellular organelle
non-membrane-bounded organelle
intracellular non-membrane-bounded organelle
neuron projection
cytoskeleton
synapse
cell projection
plasma membrane bounded cell projection
cell junction
4 5 6 7 8
Odds Ratio
-log10(adj p-value)
40 35 30 25 20
count
100 200 300
1 14 1
intracellular anatomical structure
postsynaptic density
asymmetric synapse
neuron to neuron synapse
postsynapse
organelle
cytoskeleton
neuron projection
intracellular organelle
non-membrane-bounded organelle
intracellular non-membrane-bounded organelle
synapse
cell junction
cell projection
plasma membrane bounded cell projection
4 6 8 10
Odds Ratio
-log10(adj p-value)
32 28 24
count
100 150 200 250 300
C Molecular function
Resistant vs Susceptible
Resistant vs Intermediate
Molecular Function
ATP-dependent activity
protein binding
cytoskeletal protein binding
ATP binding
adenyl ribonucleotide binding
adenyl nucleotide binding
purine ribonucleoside triphosphate binding
purine ribonucleotide binding
ribonucleotide binding
enzyme binding
purine nucleotide binding
nucleotide binding
nucleoside phosphate binding
anion binding
heterocyclic compound binding
3 4 5
Odds Ratio
-log10(adj p-value)
25.0 22.5 20.0 17.5 15.0
count
100 200 300
9 6 9
structural constituent of synapse
binding
actin filament binding
protein binding
actin binding
microtubule binding
cytoskeletal protein binding
tubulin binding
ATP-dependent activity
protein domain specific binding
molecular adaptor activity
enzyme binding
protein-containing complex binding
ATP binding
adenyl ribonucleotide binding
10 20 30
Odds Ratio
-log10(adj p-value)
30 25 20 15 10
count
100 200 300

**Supplementary Figure 2**. **Gene ontology terms enriched in upregulated differentially expressed genes for Resistant compared to Susceptible or Intermediate subgroups.** The top 15 significantly (corrected *p*-value < 0.05) enriched biological process (**A**), cellular component (**B**), molecular function (**C**) gene ontology (GO) terms are shown in bubble plots, with Venn diagrams depicting the specific overlap of these GO terms that were associated with the Susceptible (orange) or Intermediate (green) subgroups compared to Resistant rats. (**A**) 10 of the top 15 biological processes were enriched for upregulated genes in both Susceptible or Intermediate subgroups, including cellular organization and development, whereas actin cytoskeleton dynamics and neuronal function terms were specific to upregulated genes in the Intermediate subgroup, and the chromatin remodelling and microtubule-based process terms were specific to the Susceptible rats (bold terms). (**B**) Cellular components terms largely overlapped (14/15) involving synaptic functions, cellular organisation and intracellular structures, however the somatodendritic compartment term was specific to the Susceptible subgroup comparison and the neuron to neuron synapse term was specific to the Intermediate subgroup comparison (bold terms). (**C**) Upregulated genes in both subgroups are commonly associated with binding functions (bold), although the specificity of binding terms was a mix of overlapping and distinct.

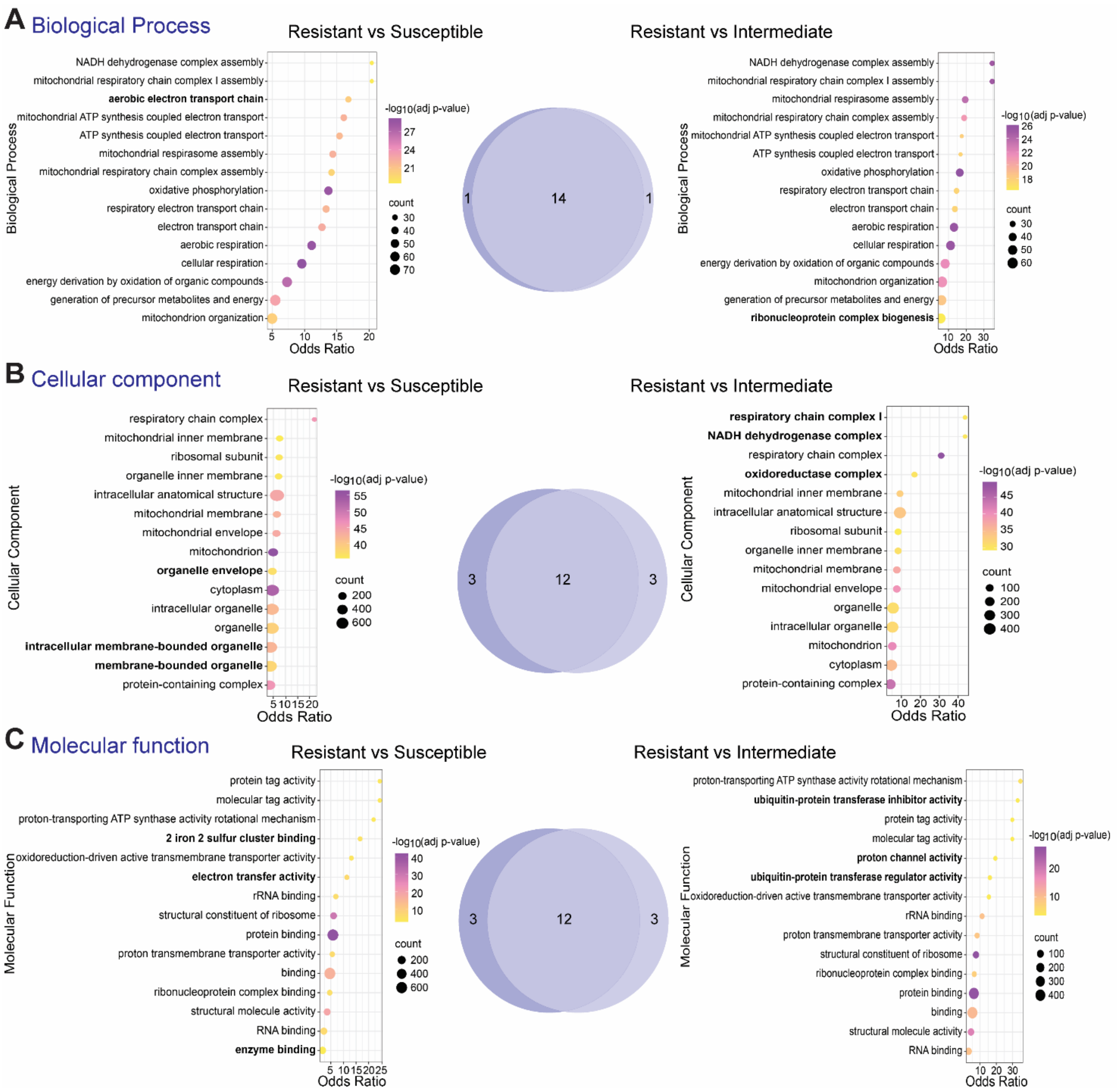


**Supplementary Figure 3. Gene ontology terms enriched in downregulated differentially expressed genes for Resistant compared to Susceptible or Intermediate subgroups.** Top 15 significantly (corrected *p*-value < 0.05) enriched biological process (**A**), cellular component (**B**), molecular function (**C**) gene ontology (GO) terms are shown in bubble plots, with Venn diagrams depicting the specific overlap of these GO terms that were associated with the Susceptible (orange) or Intermediate (green) subgroups compared to Resistant. (**A**) Biological processes mostly overlap between two subgroup comparisons (14/15), as do (**B**) the cellular components terms (12/15) and (**C**) molecular processes (12/15). The interesting differences between downregulated genes within these comparisons include (**A**) aerobic electron transport chain, (**B**) organelle envelope and (intracellular) membrane-bound organelle, (**C**) 2 iron 2 sulfur cluster binding, electron transfer activity and enzyme binding downregulated in Susceptible; and (**A**) ribonucleoprotein complex biogenesis, (**B**) respiratory chain complex, NADH dehydrogenase complex and oxidoreductase complex, (**C**)

proton channel activity, ubiquitin-protein transferase inhibitor and regulator activity downregulated in Intermediate (see bold text).
